# Unstressed cells are alike, but stressed cells differ: Environmental and single-cell heterogeneity in yeast stress responses

**DOI:** 10.1101/2025.03.12.642837

**Authors:** Rachel Eder, Leandra Brettner, Kara Schmidlin, Mohammad Donyavi, Kerry Geiler-Samerotte

## Abstract

Cellular stress responses are central to survival, adaptation, and disease, yet individual cells experiencing the same environment can differ substantially in how they respond. Understanding which features of the stress response are shared among cells, and which vary across cells and stressors, requires resolving transcriptional responses at the level of individual cells. Here, we used ultra-high-throughput single-cell RNA sequencing to profile more than 16,000 *Saccharomyces cerevisiae* cells exposed to diverse stressors, including protein misfolding, nutrient limitation, and drug stress. We compared the canonical Environmental Stress Response (ESR), originally defined from population-level measurements, with stress-responsive programs identified by emphasizing expression changes that were consistent across individual cells. Although the canonical induced ESR (iESR) showed strong average activation in stressed populations, its activation varied substantially among their constituent cells. Stress-specific programs identified by our single-cell-aware analysis were more consistent predictors of individual cell stress, yet showed little overlap with the ESR or with one another across stressors. This stress-specificity was asymmetric: Induced programs varied considerably across stressors, whereas repressed programs were dominated by a conserved set of growth- and ribosome-related genes. Consequently, unstressed cells could be identified across environments much more reliably than stressed cells, revealing a transcriptional version of the Anna Karenina principle: Unstressed cells resemble one another in their shared growth programs, whereas stressed cells diverge in stress-specific ways. Single-cell resolution also revealed an alternative transcriptional state in which cells with weaker activation of canonical stress programs showed stronger transposable element (TE) expression. Although iESR and TE expression were positively associated across stressed populations, they were inversely related across individual cells, demonstrating that population averaging can obscure, and even reverse, relationships among transcriptional programs. Together, these results show that population-level and single-cell analyses capture complementary properties of stress responses. Both perspectives are important in understanding and ultimately predicting how cell populations survive and adapt under stressful conditions.

## Introduction

Cells encounter a wide range of environmental and physiological stresses, and decades of work have established that cells mount coordinated transcriptional programs in response. In budding yeast, these studies led to the description of the Environmental Stress Response (ESR), a characteristic program marked by the induction of chaperones and other protective factors. It is also marked by the concurrent repression of growth-related processes such as ribosome biogenesis (Gasch et al. 2000; Brauer et al. 2008; Ho and Gasch 2015; Costa-Mattioli and Walter 2020; Chen et al. 2023). Originally defined from bulk transcriptomics methods, the ESR has been an invaluable framework for stress biology. This canonical response continues to serve as a barometer of stress across diverse studies, guiding interpretation of gene expression data in yeast and informing studies of stress biology in other systems (Taymaz-Nikerel et al. 2016; Gasch et al. 2017; Hackley and Schmid 2019; Costa-Mattioli and Walter 2020).

Bulk measurements aggregate expression across thousands or millions of cells, emphasizing transcriptional signals that are strong at the population level without revealing how those signals are distributed among individual cells. The advent of single-cell methods has shown that transcriptional responses to stress can be highly heterogeneous. In yeast, pioneering single-cell studies have revealed substantial cell-to-cell variability in the activation of stress-associated transcripts (Cai et al. 2008; Munsky et al. 2018; Li et al. 2018; Nadal-Ribelles et al. 2019, 2025; Sampaio et al. 2022; Bergen et al. 2022; Su et al. 2023; Kocik and Gasch 2024), while studies in cancer (He et al. 2024) and bacteria (Choudhary et al. 2023; Wang et al. 2023) suggest that heterogeneous stress responses are a general phenomenon. These observations raise a fundamental question: Are the genes that best define stress at the population level also best at defining stress in individual cells? Can a single cell’s expression level of a population-defined stress response predict whether that cell has been exposed to an external stressor? Are there other gene sets that, while not the strongest population-level indicators of stress, are more consistently activated across cells exposed to the same stress? Single-cell measurements provide an opportunity to ask which genes most reliably distinguish stressed from unstressed cells, whether those genes overlap with the canonical ESR, and whether the same genes identify stressed cells across diverse environments.

Progress in answering these questions has been limited by the scale of single-cell data available. Historically, single-cell studies of stress in yeast have either profiled only a few hundred cells, targeted a small number of genes, or focused on a single stressor. Consequently, we lack a comprehensive view of stress responses across diverse environments for large cell populations. Yeast is uniquely well suited to fill this gap because it combines rich prior knowledge of canonical stress biology (Gasch et al. 2000; Brauer et al. 2008), ultra high-throughput single-cell methods (Brettner et al. 2024), and conserved eukaryotic stress phenomena such as upregulation of heat shock proteins and transposable elements (Paquin and Williamson 1984; Lindquist and Craig 1988; Bradshaw and McEntee 1989; Horváth et al. 2017). These features make yeast not only an excellent reference model, but also a valuable system for uncovering stress mechanisms relevant to other eukaryotes.

Here, we leverage these advantages by applying an ultra high-throughput single-cell transcriptomics platform (Kuchina et al. 2021; Brettner et al. 2024) to profile approximately 16,000 yeast (*Saccharomyces cerevisiae*) cells exposed to diverse stresses, including protein misfolding, nutrient limitation, and fluconazole stress. Each cell yielded hundreds of transcripts, and by annotating transposable elements (TEs) alongside coding genes, we captured expression across more than 6,000 features. This scale provides an exceptionally rich dataset, allowing us to use analytic tools rarely accessible in microbial single-cell work. For example, we identify *de novo* stress-response programs that are obscured by population aggregates, and we study whether those programs are uniformly activated or heterogeneous (i.e., restricted to cell subpopulations). In parallel, we use the ESR as a well-established barometer of stress, providing a fixed reference to track canonical induction and repression across conditions and single cells. Together, this combination of scale, analytic power, and a trusted measure of stress allows us to probe transcriptional heterogeneity both across environments and within stressed populations.

Our high-throughput, single-cell approach reveals gene expression programs and patterns that bulk measurements miss. While we confirm that the iESR is strongly upregulated in aggregate across all stress conditions we survey, we find it is not uniformly activated across stressed cells. Instead, other stress condition-specific genes appear to be more uniform signatures of specific stresses across single cells. Even with these more uniform signatures, there appears to be an asymmetry reminiscent of the Anna Karenina principle (Zaneveld et al. 2017). Just as Tolstoy wrote that “Happy families are alike; every unhappy family is unhappy in its own way,” we find that unstressed cells resemble one another in their activation or maintenance of growth programs, whereas stressed cells are more divergent in their transcriptional programs. As a result, stress-induced genes, long used as population-level markers, perform poorly at predicting stress exposure at the single-cell level. This does not imply that canonical population-level stress responses such as the ESR are biologically uninformative. On the contrary, these responses robustly predict stress at the population level and have revealed key mechanisms of stress biology, including the activation of regulators such as Msn2/4 (Martínez-Pastor et al. 1996; Gasch et al. 2000; Causton et al. 2001). Instead, our findings highlight that bulk and single-cell analyses emphasize different aspects of cellular responses to stress.

For example, a critical insight from our single-cell analysis is that cells experiencing the same stress can partition into distinct transcriptional states, with some activating canonical stress programs and others devoting substantial transcriptional output to TEs. Bulk analyses not only obscure this heterogeneity but create the illusion that both TEs and canonical stress responses are induced simultaneously in stressed cells. In sum, our findings establish population-level and single-cell analyses as complementary perspectives: One identifies the transcriptional programs that dominate across populations, while the other reveals how unevenly those programs are distributed and what alternative states individual cells adopt.

## Results

### All stressors surveyed elicit a canonical stress response known as the iESR at the population level

The four stress conditions examined in this study include the misfolding of a gratuitous, heterologous protein (**Fig 1A**; top row, purple), limitation of glucose (**Fig 1A**; second row, blue), limitation of sucrose (**Fig 1A**; third row, red), and exposure to the antifungal drug fluconazole (**Fig 1A**; bottom row, green). Previous studies have characterized yeast responses to protein misfolding in bulk populations (Travers et al. 2000; Trotter et al. 2002; Geiler-Samerotte et al. 2011; Narayana Rao et al. 2022), whereas nutrient limitation and antifungal stress are well-established physiological perturbations that elicit broad transcriptional responses (Gasch et al. 2000, 2017; Causton et al. 2001; Brauer et al. 2008). Here, we use single-cell transcriptomics to examine how individual cells respond to each of these stresses.

**Fig 1.**
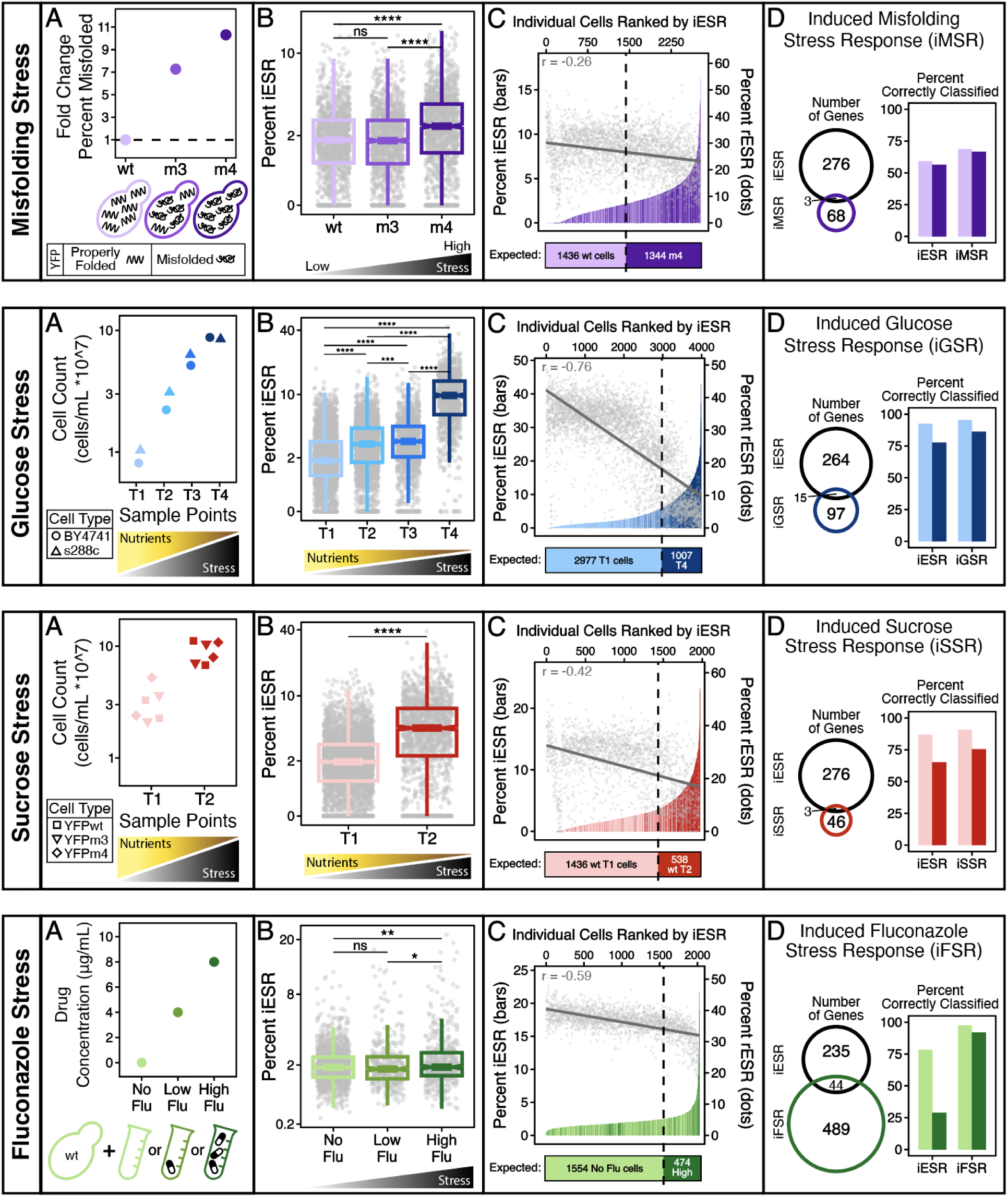
Transcriptional responses differ across stressors and individual cells. Each row represents a different stress paradigm: protein misfolding (purple), glucose limitation (blue), sucrose limitation (red), or fluconazole treatment (green), with darker colors indicating greater stress. Across rows, **(A)** shows the experimental design and stress conditions, **(B)** shows that the percent of the transcriptome devoted to the canonical induced Environmental Stress Response (iESR) generally increases with stress, **(C)** shows that iESR levels nevertheless do not cleanly distinguish stressed from unstressed individual cells, and **(D)** compares the iESR with stress-specific gene sets identified using our single-cell-aware analysis and their ability to classify cell provenance. **(A, top)** YFPm3 and YFPm4 expressing strains have 7-fold and 10-fold, respectively, more misfolded protein than YFPwt, as measured in previous work using Western blots (Geiler-Samerotte et al. 2011). **(B, top)** The percent of the transcriptome devoted to the iESR increases with misfolding stress. Each dot represents a cell sampled in early log-linear growth; boxplots display the median and quartiles, **** indicates *p* < 0.0001 with Wilcoxon rank-sum test; ns indicates no significance. **(C, top)** YFPwt and YFPm4 cells are ordered by the percent of the transcriptome dedicated to the iESR (bars). An inverse correlation is observed between the (induced) iESR (bars) and (repressed) rESR (points); the Spearman correlation (upper left) is negative; the upper-left corner reports Spearman’s ρ, and the solid line shows the linear best fit. The vertical dashed line marks the number of unstressed cells and therefore the expected classification boundary if unstressed cells have lower iESR induction. **(D, top)** Venn diagram shows only a small overlap in iESR and iMSR genes. Bars display the percent of correctly classified YFPwt or YFPm4 cells based on the rank order of their iESR or iMSR levels. **(A, row 2)** Haploid and diploid yeast strains were sampled at four timepoints during growth in glucose-limited medium; increasing cell density corresponds to increasing nutrient limitation. Cell density is plotted on a log10 scale. **(B–D, row 2)** Panels are as described above; *** indicates p < 0.001. **(A, row 3)** YFPwt, YFPm3, and YFPm4 strains were sampled at two timepoints during growth in sucrose-limited medium; increasing cell density corresponds to nutrient limitation. **(B–D, row 3)** Panels are as described above; **panel C** shows YFPwt cells only. **(A, bottom)** A single yeast strain (“wt bc”) was sampled during exponential growth at increasing fluconazole concentrations. **(B–D, bottom)** Panels are as described above; ** indicates p < 0.01 and * indicates p < 0.05.

To investigate protein misfolding stress, we utilized strains expressing increasingly misfolded variants of yellow fluorescent protein (YFP) (Geiler-Samerotte et al. 2011, 2013). We used three strains: one in which very few YFP molecules misfold (YFPwt), one in which almost every YFP molecule misfolds (YFPm4), and an intermediate strain (YFPm3) (**Fig 1A**; top row, purple) (Geiler-Samerotte et al. 2011, 2013). Because all three strains express YFP from the same smoothly inducible promoter, the only appreciable difference between them has previously been shown to be the fraction of YFP that misfolds (Geiler-Samerotte et al. 2011, 2013). For nutrient stress, we compared batch cultures sampled early in growth, when nutrients were abundant, to cells sampled later from the same flask after either glucose or sucrose had been partially depleted (**Fig 1A; Fig. S1**). For fluconazole stress, we compared untreated cells with cells grown in 4 or 8 µg/mL fluconazole.

To determine whether individual cells mount transcriptional responses to these stresses, we performed single-cell RNA-seq on each condition (Brettner et al. 2024). We quantified the fraction of each cell’s transcriptome (Hao et al. 2024) devoted to the induced Environmental Stress Response (iESR) (Gasch et al. 2000). This approach was motivated by growth-law and resource-allocation frameworks, which conceptualize transcriptomes as partitioned into functional sectors whose relative allocation changes across conditions (Scott et al. 2011; Klumpp and Hwa 2014). Focusing on transcriptome fractions rather than read counts of gene programs is common in scRNA-seq analyses (McCarthy et al. 2017; Mead et al. 2018; Pont et al. 2019), because it helps account for cell-to-cell variation in total captured RNA, sequencing depth, and cell size (Vallejos et al. 2017).

As expected, the median fraction of transcripts devoted to the iESR increased with the severity of protein misfolding, from YFPwt to YFPm4 cells (**Fig 1B**; top row). Likewise, iESR allocation increased over time as cells depleted either glucose or sucrose from the growth medium (**Fig. 1B**; rows 2 and 3) and as drug concentration increased (**Fig 1B**; bottom row). These results reinforce the utility of the iESR as a barometer of stress across diverse environments, consistent with previous studies (Gasch et al. 2000, 2017; Causton et al. 2001; Brauer et al. 2008). In all three of these stress paradigms, however, the distributions of stressed and unstressed cells overlapped substantially, with some ostensibly unstressed cells exhibiting stronger iESR induction than more severely stressed cells (**Fig 1B**; each gray point represents one cell).

### Not all stressed cells mount the canonical transcriptional response to these stresses

The overlapping distributions in **Fig 1B** suggest that, unlike in studies using stronger stressors (Gasch et al. 2017), transcriptional profiles based on the canonical iESR do not reliably distinguish stressed from unstressed cells in our study. Indeed, when we rank order cells based on the percent of their transcriptome devoted to the iESR, our stressed (**Fig 1C**; darker colors) versus unstressed (**Fig 1C**; lighter colors) cells do not separate cleanly. For example, when we attempt to classify YFP-expressing cells as either YFPwt (1436 cells) or YFPm4 (1344 cells) by ordering them based on the percent transcriptome devoted to iESR, only 59% of YFPwt and 56% of YFPm4 cells fall into the expected category (**Fig 1C** and **Fig 1D**; row 1 first set of barplots). Even fewer high fluconazole-stressed cells (only 29%) are correctly identified using this method (**Fig 1D**; bottom row). Using the same simple rank ordering procedure to identify which cells have experienced nutrient limitation results in a higher success rate (**Fig 1C**), we correctly identify 78% (glucose) and 65% (sucrose) of nutrient stressed cells (**Fig 1D**; rows 2 and 3 first set of barplots). Still, a significant fraction of cells are incorrectly classified as coming from the unstressed condition in all four experiments (**Fig 1D**; first set of barplots, darker bars never exceed 80%). These misclassified cells are not simply “borderline” cases. Several cells that experienced stress show very low iESR activation (see dark purple, red, blue or green lines at the left edges of the distributions in **Figs 1C**). These errors become even worse under less extreme versions of these stressors (see additional stress conditions in **Fig S2**) and when we report normalized read counts of iESR genes rather than the percent of the transcriptome, as the former is more sensitive to variation in cell size and sampling efficiency (**Fig S3**). Taken together, these results suggest that some cells with a history of stress are behaving alternatively to cells that do mount a canonical stress response.

These observations raise a central question: Why do many cells sampled from stressful conditions show little activation of the canonical iESR? One possibility stems from how the iESR was originally defined using population-aggregated data. Bulk analyses preferentially identify genes with large accumulated changes in expression, even when those changes are heterogeneous across individual cells. Thus, although the canonical iESR accurately describes a strong population-level response, its induction may not occur consistently enough across cells to reliably indicate the environmental history of any individual cell. A related possibility is that each stressor elicits a distinct, stress-specific transcriptional program that is more consistent across cells and therefore more informative for predicting cell provenance. To explore these possibilities, we used our single-cell data to define stress-responsive gene sets by emphasizing genes whose expression differed significantly and consistently between cells sampled from stressed and unstressed conditions.

### Each stressor provokes a distinct transcriptional response

Previous work has already identified the strongest population-level stress responses in yeast, including the ESR (Gasch et al. 2000). Here, we wanted to leverage our single-cell data to identify a different kind of stress signature: genes whose expression changes consistently enough across cells to help distinguish cells sampled from stressed versus unstressed conditions. To do this, we defined stress-responsive genes directly from our single-cell datasets. For each of the four stress types (rows of **Fig 1**), we repeatedly subsampled cells from the stressed and unstressed conditions and tested each gene for differential expression. To prevent replicate flasks with more recovered cells from contributing disproportionately within a condition, we sampled the same number of cells from each flask, using the flask with the fewest cells to set the sampling depth. We repeated this procedure 50 times using independently sampled sets of cells. The key distinction of this single-cell-aware approach was that genes were selected based on the reproducibility of their response across these independent cell subsamples, rather than simply on the magnitude of their average response. We therefore classified genes as stress-induced only if they were significantly upregulated after multiple-testing correction in at least 75% of the subsampling iterations and showed a consistent direction of change.

This approach differs from traditional bulk analyses, where stress-responsive genes are often identified from aggregate expression changes across a small number of replicate populations. Bulk analyses are well suited to detecting strong average responses, even when those responses are driven by only a subset of cells. By contrast, our single-cell-aware approach emphasizes expression changes that are reproducible across different subsamples. Thus, rather than asking which genes show the largest average response to stress, we asked which genes most consistently distinguish stressed from unstressed single cells.

In each of the four stress conditions, we compared gene expression between cells exposed to the highest and lowest levels of each stressor (**Fig 1; Fig S4**). For each stress type, this analysis identified a distinct set of genes with significantly elevated expression under the most severe condition relative to the least severe condition (**Supplemental Table 1; Fig S5**). We refer to these programs as the induced misfolding stress response (iMSR), induced glucose stress response (iGSR), induced sucrose stress response (iSSR), and induced fluconazole stress response (iFSR). Gene ontology analysis showed that all four programs are enriched for stress-responsive functions (**Supplemental Table 2**), yet each exhibits surprisingly little overlap with the canonical iESR (**Fig. 1D**; Venn diagrams). Likewise, the four stress-specific programs are largely distinct from one another. Across all four, only seven genes are shared, including the Ty1 and Ty2 transposable-element families, which were represented at the family level in this analysis (**Fig S6; Supplemental Table 1**). Thus, when stress-responsive programs are defined using analytical approaches that emphasize consistent expression changes across single cells, the resulting induced gene sets appear more stress-specific.

Notably, activation of these stress programs is better at predicting which cells were exposed to stress than the iESR. A similar threshold classifier to that in **Fig 1C**, but where cells are rank ordered based on the percent of their transcriptome devoted to these stress responses (**Fig S7**), correctly classifies more cells than the iESR (**Fig 1D**). The iFSR provides the clearest example of the utility of these stress-specific programs for predicting cell provenance. A threshold classifier that uses iFSR gene levels predicts stressed cells with over 90% accuracy, while one using iESR gene levels correctly identifies less than 30% of cells exposed to fluconazole stress (**Fig 1D**; bottom row; **Fig. S7**). The same pattern extends to the remaining most severe stress conditions (**Fig 1D**) and most lesser stresses (**Fig S2**), where the iMSR, iGSR, and iSSR each classify stressed cells more accurately than the canonical iESR (**Fig S7**).

The differences between the stress-specific gene sets we identify and the canonical iESR could be biological: Our experiments examine different and often milder, sustained stresses than the acute perturbations originally used to define the iESR. However, it could also reflect analytical differences between population-level and single-cell-aware approaches. We next examine this possibility by comparing the average magnitude and cell-to-cell consistency of the expression changes captured by each response.

### Population versus single-cell approaches reveal different views of stress responses

Single-cell data provide new ways to measure differences in gene expression across environments. Traditional bulk analyses average gene expression across many cells. They may favor detection of genes with strong responses to environmental change that are not necessarily uniform across all cells exposed to a given environment. By contrast, in single-cell analyses, each cell can serve as an independent observation, which can increase power to detect subtle changes when they occur consistently across many cells. This means the two approaches may be biased in different ways. Population-level methods may highlight genes with strong changes that may be driven by only a subset of cells, whereas our single-cell-aware method may preferentially detect smaller but consistent shifts across many cells. Both types of signals are important.

To show that population-level and single-cell-aware approaches can favor the detection of different gene sets, we compared two properties of each gene’s response to stress: (1) the average log2 fold change in expression between stressed and unstressed cells (**Fig 2A**), and (2) the consistency of that fold change across independently subsampled groups of cells (**Fig 2B**). For each stress comparison, we generated 50 independently balanced subsamples of stressed and unstressed cells and calculated a separate fold change for every gene in each subsample. **Fig 2A** shows each gene’s average fold change across the 50 subsampling iterations, whereas **Fig 2B** shows the coefficient of variation in its fold change across those same iterations. The average fold change of iESR genes is significantly positive in all stressed relative to unstressed conditions compared to all other genes (**Fig 2A**; black points). This is consistent with these genes being important markers of stress at the population level. The expression levels of the stress-responsive gene sets that we detected using our single-cell-aware approach (iMSR, iGSR, iSSR, and iFSR) are elevated as well, though not always as much as the iESR (**Fig 2A**; colored points). This pattern changes when we look at the variation in gene expression (**Fig 2B**). The gene sets that we defined have significantly lower variation in gene expression across subsets of cells. This is consistent with the idea that single-cell-aware analysis can detect gene sets that are more uniformly responsive to a given stress, while population analyses preferentially detect gene sets with strong average responses that are not necessarily predictive of single-cell state. Together, these analyses show that the discrepancy between the canonical ESR and the stress-specific responses identified here is not simply a failure to induce the ESR. Instead, population averaging and single-cell-aware analysis prioritize different statistical properties of the response: the magnitude of the average expression change and the consistency of that change across individual cells, respectively.

**Fig 2:**
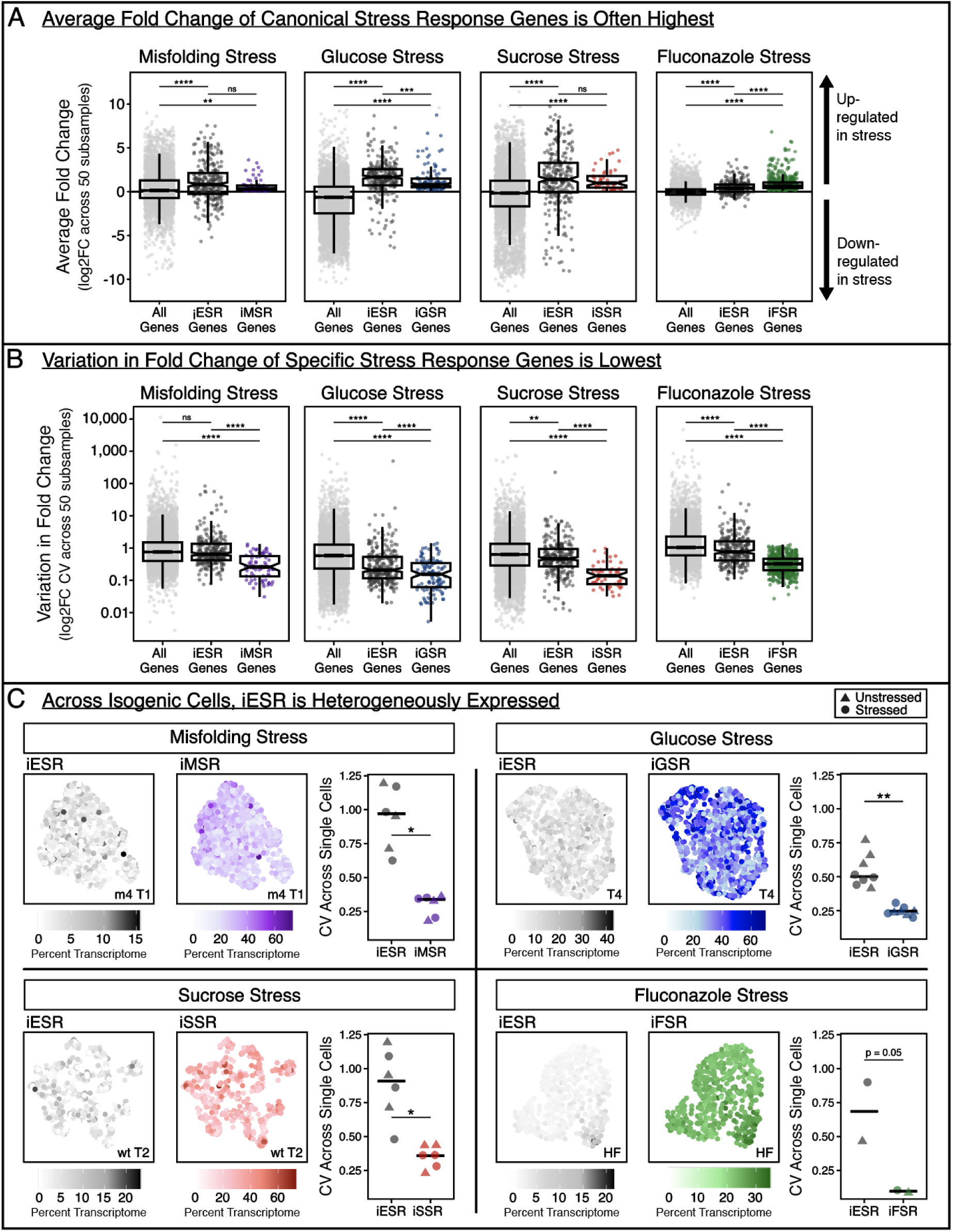
While iESR genes tend to be upregulated in stressful conditions, their expression is heterogeneous. **(A)** Vertical axis displays the average log2 fold change (FC) between stressed and unstressed cells across 50 independently balanced subsampling iterations. Each point represents FC for a single gene. Black points correspond to the iESR genes (**Supplemental Table 1**), colored points to the genes that define each stress-specific response (**Supplemental Table 1**), and gray points to all other genes. A positive average FC corresponds with genes upregulated in stressed cells. Boxplots show the median and quartiles across all genes in each category, notches on boxplots represent 95% confidence intervals, **** indicates *p* < 0.0001 with Wilcoxon rank-sum test; *** indicates *p* < 0.001; ** indicates *p* < 0.01, and ns indicates no significance. **(B)** Coefficient of variation in log2 fold change across the same 50 independently balanced stressed-versus-unstressed cell subsamples. Lower values indicate genes whose response to stress was more consistent across different samples of single cells. **(C)** Every point in each UMAP represents a single cell sampled from one of the four stress conditions: misfolding (top left, YFPm4 cells), glucose limitation (top right, timepoint 4 cells), sucrose limitation (bottom left, timepoint 2 cells), and fluconazole stress (bottom right, 8 ug/uL fluconazole cells). The pair of UMAPs in each quadrant are identical except for the way the cells are colored. The leftmost UMAP in each pair is colored based on the percent of each cell’s transcriptome devoted to the iESR. The rightmost UMAP in each pair is colored based on the percent of each cell’s transcriptome devoted to the stress-specific response defined in Fig 1. All replicate experiments corresponding to the condition indicated on each UMAP were merged, integrated, and normalized prior to plotting. Since this panel depicts percent expression, as opposed to fold change in expression, the average iESR signal appears weaker than in **panel A**. The salient point is that there is substantial heterogeneity in iESR induction from cell to cell. This is similarly shown in the graph which plots the coefficient of variation (CV) of percent expression of each gene set across single cells in every replicate flask, including the unstressed conditions, with horizontal bars depicting the median across flasks. CVs were compared using paired two-sided Wilcoxon signed-rank tests, as each replicate provided matched measurements of iESR and iXSR variability. * indicates *p* < 0.05 and ** indicates *p* < 0.01. The CV for iESR induction is always greater than for the other stress responses.

An independent population-level dataset further supports the conclusion that analytical approach influences which gene sets are recovered. Brauer et al. measured gene expression using bulk DNA microarrays under six nutrient limitations, including glucose, and identified genes whose expression was significantly correlated with growth rate (Brauer et al. 2008). Among genes represented in their dataset, 64.7% of the iESR genes changed significantly across nutrient levels, compared with 39.3% of the iGSR defined here. This pattern supports the idea that the analytical method can be a major determinant of which stress-responsive genes are identified: Population-level analyses repeatedly emphasize iESR genes, whereas our single-cell-aware analysis emphasizes genes whose responses are more consistent across individual cells.

Within our four stress conditions, one stress deviates from the general pattern: fluconazole stress. In the three non-drug stresses, the canonical iESR remains among the most strongly upregulated gene set at the population level (**Fig 2A**). Thus, the discrepancy between the stress-response genes we detected and the canonical iESR is not necessarily due to the fact that our experiments involve continuous mild stress rather than the short-term acute stresses originally used to define the iESR (Gasch et al. 2000). Instead, the discrepancy appears to be largely methodological. Even in most of our continuous, subtle stress conditions, the iESR captures genes with the strongest average induction across populations (**Fig 2A**), whereas our single-cell approach identifies genes whose expression changes most consistently across individual cells. Fluconazole produces a different result. The iFSR genes are more strongly induced than the canonical iESR even at the population level (**Fig 2A**), suggesting that the limited overlap in **Fig 1D** between the iESR and iFSR genes reflects not only differences in analytical approach but also distinct biology of the transcriptional response to 8 µg/mL fluconazole under our experimental conditions.

We obtain the same general result when we examine expression variation across stressed cells (**Fig 2C**) rather than variation in fold changes across subsets between stressed and unstressed conditions (**Fig 2B**). iESR induction in stressed cells appears more heterogeneous, with some stressed cells devoting very little of their transcriptome to the response (**Fig 2C**; gray UMAPs). Note that here we are looking at only stressed cells, and not at fold change in stressed versus unstressed conditions, which is why here the average iESR signal appears weaker than the stress-specific signals. In contrast to iESR induction, the stress-specific responses appear more uniformly expressed, with a larger fraction of stressed cells exhibiting moderate induction (**Fig. 2C**; colored UMAPs). We can quantify variation across each group of cells to formally test this observation. The coefficient of variation for iESR induction across cells is always significantly higher than that of the corresponding stress-specific response (iMSR, iGSR, iSSR, or iFSR; **Fig 2C**, boxplots). Thus, analyses across cells (**Fig 2C**) and across genes (**Fig 2A–B**) lead to the same conclusion: Gene sets identified from population-level studies, such as the iESR, emphasize average fold changes in expression between populations, whereas our single-cell-aware approach preferentially identifies consistent shifts in expression across cells.

### At the single-cell resolution, stress-repressed genes generalize better across stressors than stress-induced genes

Our analyses thus far suggest that, when defined using our single-cell aware approach, the transcriptional programs induced by different stresses are largely distinct. This observation led us to ask whether, at the single-cell level, stress responses follow a broader organizing principle. Specifically, if each stress induces its own characteristic transcriptional program whereas unstressed cells maintain a common growth-oriented state, then stressed cells should resemble one another less than unstressed cells. This idea parallels the Anna Karenina principle (Zaneveld et al. 2017): Healthy systems are similar because they satisfy many shared requirements, whereas perturbed systems can fail in many different ways.

To test this idea, we compared both the induced and repressed transcriptional programs identified for each stress condition. The induced responses (iESR, iMSR, iGSR, iSSR, and iFSR) were remarkably stress-specific, with most genes appearing in only a single stress response (**Fig 3A**; black bars). In contrast, the stress-repressed responses (rESR, rMSR, rGSR, rSSR, and rFSR) shared many more genes across conditions (i.e., the fraction of genes that are unique is higher for corresponding stress-induced than stress-repressed genes) (**Fig 3A**; yellow bars). These repressed gene sets are enriched for growth-related functions, including ribosome biogenesis, consistent with previous descriptions of the rESR (Gasch et al. 2000; Brauer et al. 2008), and their expression anticorrelates with the induced responses (**Figs 1C** and **S2**). Thus, while stressed cells induce distinct, condition-specific transcriptional programs, they repress a much more conserved set of growth-related genes.

**Fig 3:**
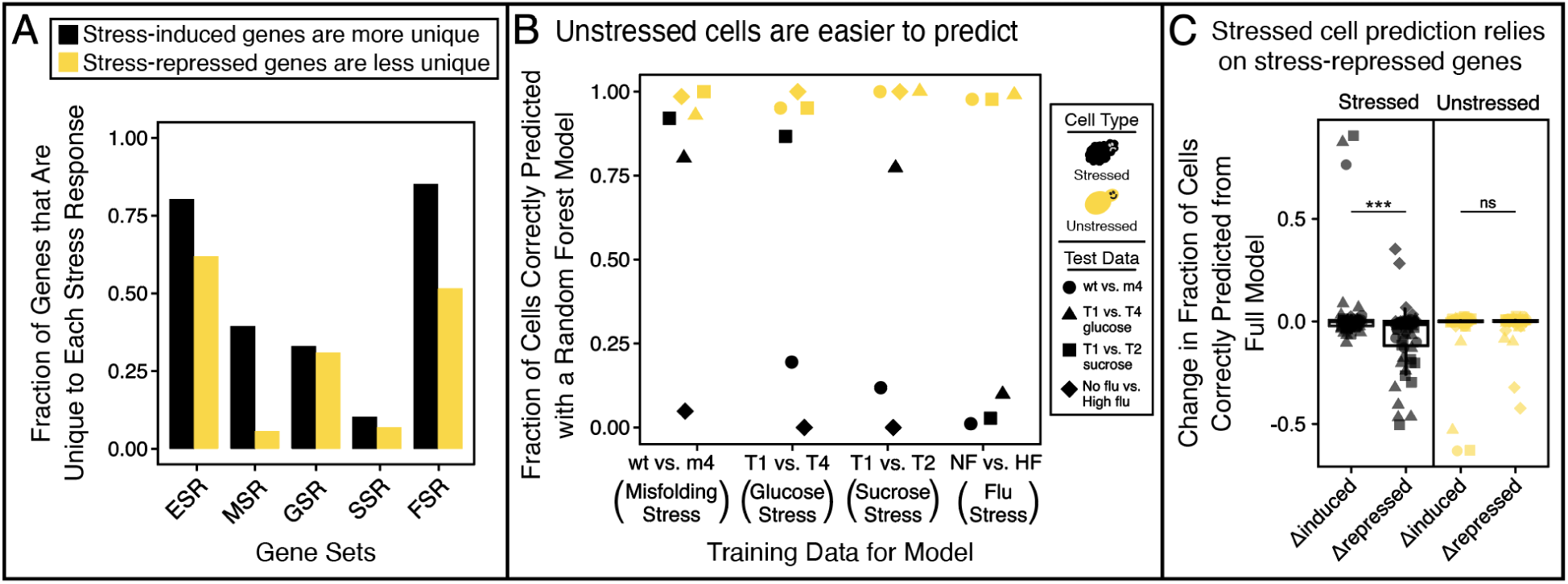
Stress-induced genes vary across stressors, making it challenging to learn a generalizable signature of stress. **(A)** Bars show the fraction of genes unique to each induced (black) or repressed (yellow) stress response across the ESR, MSR, GSR, SSR, and FSR. Genes were considered unique if they appeared in only one response. Stress-repressed gene sets are consistently less unique than stress-induced gene sets. **(B)** A Random Forest (Breiman 2001) model can correctly predict which cells are unstressed (yellow) more often than which cells have experienced stress (black). Categories on the horizontal axis correspond to the training data used to build a predictive model. The fraction of the test cells from other stress conditions correctly predicted are plotted on the vertical axis. The shape of the points corresponds to the cells used in the test data, as indicated by the legend. **(C)** Withholding stress-repressed genes significantly reduced prediction accuracy for stressed cells compared with withholding stress-induced genes (two-sided Wilcoxon signed-rank test; *** indicates *p* < 0.001), whereas prediction of unstressed cells was unaffected (ns). Models in **B** were retrained after withholding each induced or repressed stress-response gene set individually, and then they were used to predict cells from the other stress conditions; plotted values show the change in accuracy relative to the corresponding full model.

If unstressed cells are indeed transcriptionally more alike than stressed cells, then they should also be easier to recognize across datasets. To test this prediction, we trained Random Forest classifiers (Breiman 2001) on one stress experiment and asked whether they could identify stressed and unstressed cells from a different stress experiment. These models were intended as predictive tools rather than mechanistic models, allowing us to ask which transcriptional signatures generalize across diverse perturbations. Across every training-testing combination, unstressed cells were identified with greater than 90% accuracy (**Fig 3B**; yellow points), whereas stressed cells were classified much less reliably, with prediction accuracy falling below 25% in some comparisons (**Fig 3B**; black points). This asymmetry mirrors the pattern observed with our simpler threshold classifier (**Fig 1D**), where unstressed cells were also classified more accurately than stressed cells, and this trend persists across additional datasets and stress magnitudes (**Figs S4C** and **S7C**), although classification of unstressed cells becomes even more accurate under more severe stress (**Fig S4**). Together, these results support the Anna Karenina view that unstressed cells share a common transcriptional state, whereas stressed cells diverge according to the specific challenges they encounter.

We next asked which transcriptional programs account for this difference in predictability. Removing the stress-induced gene sets from the Random Forest models had little effect on prediction accuracy (**Fig 3C**). In contrast, removing the stress-repressed genes significantly reduced the ability to identify stressed cells across datasets, while having little effect on identification of unstressed cells (**Fig 3C**). Thus, successful cross-condition prediction depends primarily on the conserved repression of growth-related genes rather than on the diverse stress-specific induction programs. In other words, cells are easiest to recognize as unstressed because they all maintain a similar growth program, whereas recognizing which cells are stressed requires transcriptional programs that differ from one stress to the next.

Taken together, these findings support a model in which transcriptional stress responses comprise a conserved repression module, reflecting reduced investment in growth-related processes, together with flexible induction modules that reflect the specific nature of each stressor (Bergen et al. 2022). They do not redefine or diminish the canonical iESR, which remains a robust marker of population-level stress. Rather, they show that single-cell analyses reveal an additional layer of organization: Unstressed cells are transcriptionally more alike, whereas stressed cells are unique in ways that depend on the stress they experience.

### Cells experiencing the same stress can upregulate distinct transcriptional programs

The analyses above show that iESR activation varies substantially among individual cells and can remain very low even in cells sampled from stressful conditions (**Fig 1C**; **Fig 2B–C**; **Fig 3**). This raises a question: What are these low-iESR cells expressing instead? To address this, we pooled cells sampled from the four stressful conditions and compared gene expression between cells in the highest and lowest quartiles of iESR activation (**Fig 4A**). As expected from the inverse relationship between the induced and repressed ESR observed above (**Fig 1C**), many genes enriched in low-iESR cells were constituents of the rESR. More unexpectedly, transposable elements (TEs) were also among the transcripts enriched in low-iESR cells sampled from stressful conditions (**Fig 4A**). The increased TE signal included loci from multiple Ty families, and Ty1 reads mapped to both the 5′ and 3′ regions (**Fig S10**), suggesting the increased TE signal was not limited to an internally initiated Ty1 transcript that produces the retrotransposition inhibitor p22 (Saha et al. 2015; Ahn et al. 2017).

**Fig 4:**
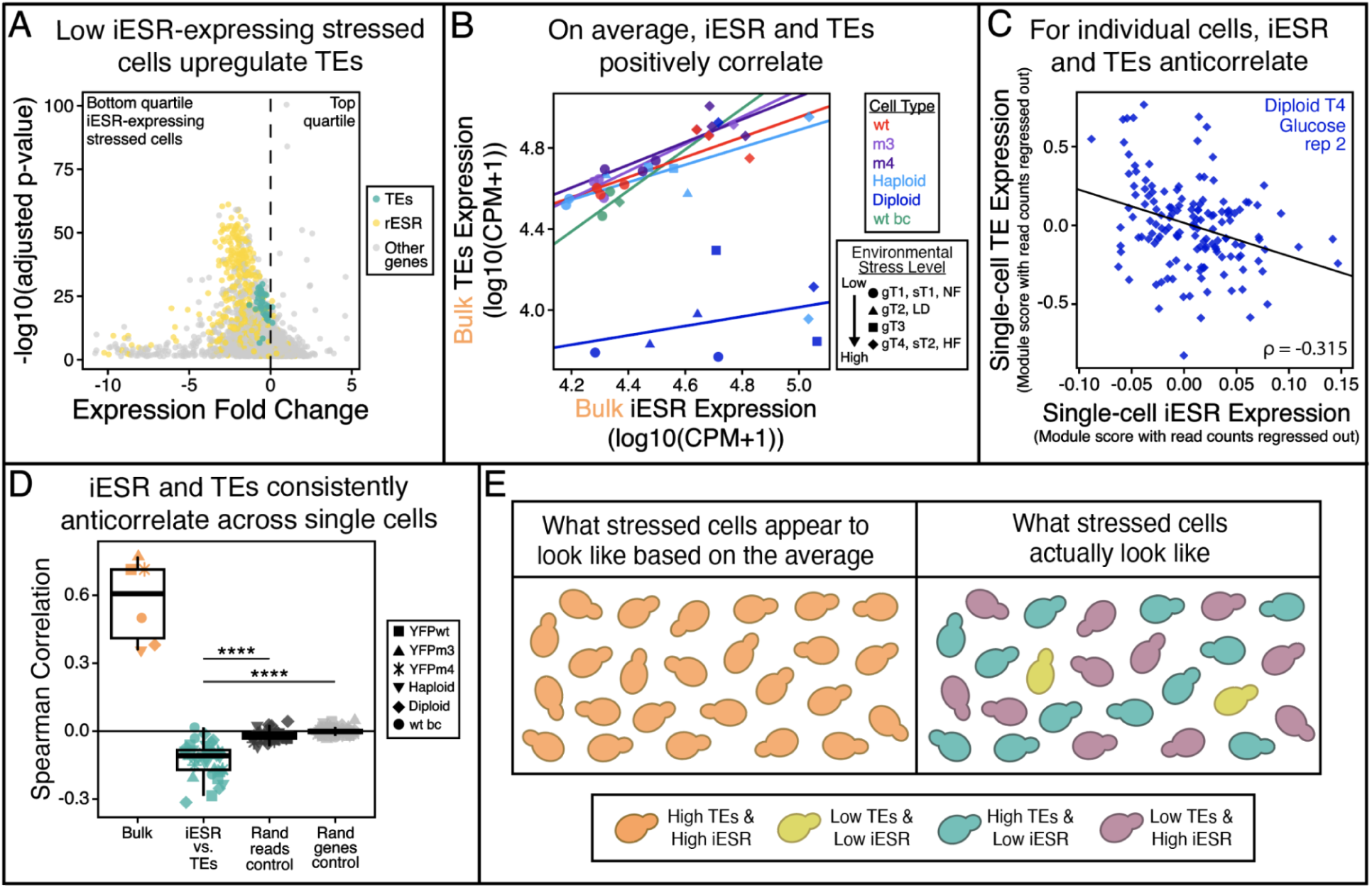
TE and iESR expression are positively associated across cell populations but negatively associated across individual cells. **(A)** Volcano plot comparing gene expression between cells with high versus low iESR activation. Cells from the four strongest stress conditions (YFPm4, glucose T4, sucrose T2, and high fluconazole) were pooled and divided into the highest and lowest quartiles of iESR activation. Genes enriched in low-iESR cells are plotted to the left and include components of the rESR (yellow) and transposable elements (TEs; teal). **(B)** At the population level, iESR and TE expression increase together. Each point represents one experimental condition (37 total), with raw transcript counts from all cells in that condition pooled to generate a pseudobulk expression value. iESR and Ty expression are shown as log10 counts per million plus 1 (log10(CPM + 1)). Lines show linear fits across conditions corresponding to increasing levels of the same stress. One outlier in each of the glucose diploid and haploid series was excluded from the corresponding fitted line but remains plotted and was retained for Spearman correlation analyses. Spearman correlations for these population-level comparisons are summarized in **panel D** (“bulk”). **(C)** At the single-cell level, the relationship is reversed. Shown is one representative sample, S288C cells at T4 of glucose limitation, in which each point represents an individual cell. iESR and TE expression were quantified using Seurat module scores; sequencing depth was regressed out when it explained at least 10% of module-score variation. Higher TE expression is associated with lower iESR activation (Spearman correlation = −0.315). Corresponding analyses for all samples are shown in **Fig S8** and summarized in **panel D**. **(D)** Comparison of population-level and single-cell relationships between TE and iESR expression and two null controls. Orange points show Spearman correlations calculated across cell populations experiencing different levels of the same stress (“bulk”); these correlations are wholly positive. Teal points show Spearman correlations calculated across individual cells within each of the 37 samples; these correlations tend to be negative. Dark gray points show random-read controls that estimate correlations expected from transcript sampling alone, and light gray points show random-gene controls that estimate correlations expected from arbitrary gene-set comparisons. The observed single-cell TE–iESR correlations are significantly more negative than either control. Statistical significance was assessed using two-sided Wilcoxon signed-rank tests; **** indicates *p* < 0.0001. **(E)** Schematic illustrating how population averaging can obscure the single-cell relationship between TE and iESR expression. Because both programs increase on average across stressed populations (**panel B**), population-level measurements could suggest that individual stressed cells activate both programs simultaneously. Single-cell measurements instead reveal that TE and iESR expression are inversely related, with cells tending toward higher expression of one program or the other (**panels C–E**).

Previous work has shown that TE expression can increase in response to environmental stress and nutrient limitation (Paquin and Williamson 1984; Rolfe et al. 1986; Bradshaw and McEntee 1989; Stanley et al. 2010; Servant et al. 2012; Curcio et al. 2015). Our data reveal an additional feature of this response: Increased Ty expression is not distributed uniformly across stressed cells but is concentrated among cells with relatively weak activation of the canonical iESR. This heterogeneity is not apparent when expression is averaged across cells. When each cell population is represented by a bulk expression value, TE expression and iESR activation are positively correlated (**Fig 4B**). This correlation could be misinterpreted to mean that individual stressed cells tend to activate the iESR and TEs together. But the single-cell data reveal the opposite relationship (**Fig 4C**). Within nearly every cell population examined in our study (**Fig S8**), cells with higher TE expression tend to have lower iESR activation, whereas cells with stronger iESR activation tend to express fewer TE transcripts (**Fig 4D**). This inverse relationship is observed across different strain backgrounds, stressors, and stress magnitudes. It also extends beyond the canonical iESR: TE expression is negatively correlated with the non-TE components of the iMSR, iGSR, iSSR and iFSR across individual cells (**Fig S9**).

One potential concern is that the negative relationship between TE and iESR expression could arise partly from how transcriptome fractions are calculated. Because each gene set is measured as a fraction of a cell’s total captured transcripts, a large increase in abundant TE transcripts could mechanically reduce the fraction assigned to the iESR even if iESR transcript abundance itself did not decrease. This is a general consequence of compositional transcriptome data (Quinn et al. 2018). We therefore conducted the above analysis using Seurat module scores, which compare the average log-normalized expression of a target gene set with that of expression-matched control genes (Tirosh et al. 2016; Hao et al. 2024). Critically, this method does not measure both programs using a shared total-transcript denominator. Where module scores retained an association with sequencing depth, we regressed out read count. The negative relationship between TE and iESR expression activity remained evident using this method (**Fig 4D**), indicating that the relationship is not explained solely by the compositional nature of transcriptome fractions.

To further test whether this negative relationship reflects specific biological structure rather than a generic artifact of transcript sampling or comparing two gene sets, we performed two complementary controls. For the first control, we bioinformatically pooled the reads for each sample to form an expression probability distribution. We then randomly redistributed transcripts to control cells, keeping the read count structure of the original cells. We then recalculated the iESR and TE module scores and the Spearman correlations for each sample 100 times and report the median as the random read controls (**Fig 4D**; dark gray). The second control consists of comparing the iESR or TE module scores to expression scores of randomly selected gene sets of the same size of the iESR and TE gene sets 100 times for each sample. The median random gene control Spearman correlation is reported for each sample in light gray (**Fig 4D**). The true correlations are significantly more negative than either control, suggesting they result from biologically relevant structure within the true data.

The connection between TEs and stress responsive genes is intriguing, because TE activation and stress response dysregulation have both separately been implicated in aging, neurodegeneration, and protein misfolding diseases, including Alzheimer’s disease (Wood and Helfand 2013; Feng et al. 2025). Elevated TE expression could represent a failure to maintain normal TE regulation or could also represent a feature of physiological states that differ in growth, replicative age, or progression toward stationary phase (Aragon et al. 2008; Patterson et al. 2015; Horváth et al. 2017; Peifer and Maxwell 2018; Specchia and Bozzetti 2021). Our data do not establish why cells adopt these different transcriptional states. Instead, our results establish the underlying pattern: Cells sampled from the same stressful environments can pursue distinct transcriptional programs, with some strongly activating the canonical iESR and others expressing high levels of Ty1 and Ty2.

This result illustrates a central distinction between population-level and single-cell analyses. Population aggregates show that both the iESR and TEs increase during stress, but they can not reveal whether those increases occur within the same cells. At single-cell resolution, the two programs are inversely related and largely concentrated in different subsets of the population. Thus, averaging does not merely conceal the uneven distribution of these responses; it produces an apparent positive association that reverses the relationship observed among individual cells (**Fig 4E**).

## Discussion

Building on prior single-cell studies demonstrating stress-response heterogeneity in yeast (Gasch et al. 2017; Saint et al. 2019; Nadal-Ribelles et al. 2019, 2025; Su et al. 2023), our study uncovers additional dimensions of this heterogeneity that bulk methods overlook or sometimes misrepresent. By profiling over 16,000 yeast cells across diverse stresses, we reveal environmental and cell-to-cell heterogeneity in stress responses.

We found that stress-induced transcriptional programs are not reliable predictors of whether a cell has experienced a stressful environment. Even the canonical iESR, which is detectably upregulated across all stresses, does not reliably encode a single cell’s environmental history. By contrast, stress-repressed transcriptional programs are more reliable. Genes tied to growth and ribosome biogenesis, whose repression is a well-established feature of the yeast stress response (Gasch et al. 2000; Brauer et al. 2008; Bergen et al. 2022; Kocik and Gasch 2024), are consistently downregulated across the stressors we examined, and in our analyses this repression strongly supports prediction of a single cell’s provenance. Consistent with prior work documenting cell-to-cell variation in stress-program activation (Cai et al. 2008; Gasch et al. 2017; Li et al. 2018; Bergen et al. 2022; Nadal-Ribelles et al. 2025), we find that cells vary in the stress-associated programs they activate, but show a more conserved repression of growth-related programs when challenged. Our cross-dataset classifier models support this view, reliably identifying unstressed cells but struggling to predict stressed ones. This distinction raises a broader question: How does transcriptional heterogeneity relate not merely to a cell’s history of stress exposure but to its subsequent fate?

An important distinction is that predicting whether a cell has experienced stress is not the same as predicting what will happen to that cell under stress. Previous single-cell studies suggest that heterogeneous stress-associated states can have consequences for subsequent survival and recovery. In unstressed yeast populations, slow-growing cells with high expression of the stress-associated trehalose regulator Tsl1 are disproportionately likely to survive subsequent severe heat stress, a phenomenon proposed to represent a form of bet hedging (Levy et al. 2012). Related work has shown that pre-stress cellular state and activation of the Dot6-mediated repression program can predict recovery from osmotic stress, with stronger repression of growth-related transcripts associated with faster acclimation (Bergen et al. 2022). More recently, heterogeneous use of osmostress-responsive transcriptional programs has also been linked to differences in cellular fitness (Nadal-Ribelles et al. 2025). These studies suggest a broader significance for the heterogeneity we observe. If different transcriptional states carry different probabilities of surviving particular stresses, then understanding the distribution of states across individual cells may ultimately be necessary not only to predict the fate of a cell but also the persistence of a population in which survival can depend on only a subset of its members.

While prior studies have shown that stressed yeast populations can occupy heterogeneous regulatory and transcriptional states (Gasch et al. 2017; Saint et al. 2019; Bergen et al. 2022; Nadal-Ribelles et al. 2025), our study leverages a variety of approaches to understand how averaging across such states can mask important signals. For example, we show that subsets of cells mount expected stress programs, while other cells devote larger fractions of their transcriptome to transposable elements (TEs). At the population level, these responses to stress appear positively correlated, yet cell-level analyses reveal them to be anticorrelated such that cells are largely mounting one or the other. Bulk data not only hide this negative correlation but even invert it, underscoring how averaging can distort the biology it seeks to summarize. Methodological comparisons further reinforce this message. Bulk fold-change analysis highlights the iESR as a reliable indicator of stressed cell populations, whereas single-cell-aware significance tests and predictive models show that iESR genes vary across individual cells and are not especially informative for prediction.

More broadly, our analysis uses single-cell information not only to characterize heterogeneity within previously defined stress-response programs but also to ask how that heterogeneity should influence the definition of a stress response itself. By repeatedly subsampling cells and prioritizing genes whose differential expression is reproducible across independently sampled subsets, we identify stress-responsive programs based on cell-level consistency rather than solely on the magnitude of their average population-level response. These programs differ substantially from the canonical ESR, are generally more uniformly expressed across individual cells, and more accurately identify the environmental provenance of single cells. Thus, single-cell data can do more than reveal heterogeneity hidden within established gene sets: They can change which genes emerge as representative signatures of a cellular response.

Taken together, our results and other single-cell studies of yeast stress responses (Gasch et al. 2017; Bergen et al. 2022; Nadal-Ribelles et al. 2025) suggest that a stressed population is better understood as a distribution of cellular states than as a single average response. Our findings add another dimension to this picture by showing that accounting for this distribution can change which genes emerge as characteristic signatures of stress. Some features, particularly repression of growth-related programs, are shared broadly across cells and environments, whereas induced transcriptional responses are more heterogeneous and stress-specific. Other responses, such as canonical stress programs and TE expression, can even be partitioned among different cells despite appearing positively associated at the population level. This distinction may be particularly important when heterogeneous states differ in their ability to withstand or recover from stress: In such cases, the fate of a population may depend not on the response of an average cell but on the distribution of responses among its members. Connecting these transcriptional states to subsequent survival will therefore be an important next step toward predicting how heterogeneous cell populations persist and adapt in stressful environments.

## Methods

### Yeast strains

*Saccharomyces cerevisiae* strain *MATa ura3Δ0 his3Δ1 met17Δ0 PACT1-GAL3::SpHIS5 gal1Δ gal10Δ::LEU2 leu2Δ0::PGAL1-YFP-KanMX6* with wt, m3, and m4 YFP mutations was used for intrinsic misfolding and extrinsic nutrient limitation stress experiments in sucrose/raffinose-limited media (Geiler-Samerotte et al. 2011). These strains express YFP from a modified galactose-inducible promoter system engineered to produce smooth, unimodal induction across cells rather than the bistable/bimodal induction characteristic of the native GAL pathway. Specifically, the strains constitutively express GAL3 and lack the endogenous GAL1 and GAL10 genes, preventing cells from metabolizing galactose while reducing positive-feedback-driven bimodality in GAL induction. We also used a diploid s288c and a haploid BY4741 for glucose nutrient limitation experiments. *Saccharomyces cerevisiae* strain *MATɑ, ura3Δ0, ybr209w::Gal-Cre-KanMX-1/2URA3-loxP-Barcode-1/2URA3-HygMX-lox66/71* (abbreviated wt barcode) was used for fluconazole stress experiments (Schmidlin et al. 2024).

### Cell growth and sampling

Cells were streaked from −80°C glycerol stocks onto YPD agar and grown for 48 hours at 30°C. Single colonies were inoculated into synthetic complete medium containing either 2% glucose (glucose-limitation experiments) or 2.5% sucrose, 1.25% raffinose, and 0.625% galactose (protein-misfolding and sucrose-limitation experiments) and grown overnight with shaking at 30°C. Overnight cultures were diluted into fresh medium to approximately 300 cells/mL (15,000 cells in 50 mL) to initiate growth experiments. Cell density was monitored over time using a Coulter counter for the YFPwt, s288c, and BY4741 cells and a spectrophotometer for the wt barcode cells, and samples were collected at the indicated time points or densities, fixed, and processed for single-cell RNA sequencing as previously described (Brettner et al. 2024). Media composition, biological replication, and sampling time points differed among stress conditions as described below.

### Protein misfolding

Three YFP strains (YFPwt, YFPm3, and YFPm4) were analyzed in two independent experiments performed on separate days, yielding three biological replicates per strain (one replicate in the first experiment and two in the second). Following overnight growth, cultures were diluted into synthetic complete medium lacking glucose and containing 2.5% sucrose, 1.25% raffinose, and 0.625% galactose to induce YFP expression (Geiler-Samerotte et al. 2011). After at least 16 hours of exponential growth, cells were sampled in mid-log phase for scRNA-seq. Cell densities at the time of sampling are shown in **Fig 1A** and **S1**. When multiple replicate flasks were sampled on the same day, average cell counts are reported in **Fig 1A**.

### Glucose limitation

Two biological replicates each of diploid S288C and haploid BY4741 cells (Brettner et al. 2024) were grown in synthetic complete medium containing 2% glucose and sampled at four time points during batch growth. Cell densities at each sampling point are shown in **Fig 1A** and **S1**. Average cell counts for replicate diploid and haploid flasks at each timepoint are reported in **Fig 1A**.

### Sucrose limitation

The same cultures used for the protein-misfolding experiments were allowed to continue growing to early stationary phase, when sucrose had become limiting. Cells were sampled for scRNA-seq, with cell densities reported in **Fig 1A** and **S1**. Because these cultures simultaneously experience YFP expression and nutrient depletion, only each YFPwt replicate was compared to itself between mid-log and early stationary phase to isolate the transcriptional response to sucrose limitation.

### Fluconazole treatment

Wt barcode cells were streaked from glycerol stocks onto YPD agar and grown for 48 hours at 30°C. A single colony was inoculated into 5 mL M3 medium containing 2% glucose and grown overnight. Cultures were then diluted into 30 mL M3 medium containing 4 or 8 µg/mL fluconazole or 0.5% DMSO (vehicle control) and grown overnight at 30°C. Cells were passaged once into fresh medium containing the same drug concentration and harvested at OD600 ≈ 0.5. Cultures were divided into five 1-mL aliquots, pelleted by centrifugation, stored at −80°C, and one aliquot was used for single-cell library preparation.

### Single-cell RNA sequencing

For the YFP, s288c, and BY4741 cells, 3mL of liquid culture were removed from the shaking flasks at each timepoint. From that, 100uL were used to count cells (as described above). The remaining 2.9mL of liquid culture was immediately fixed with 390uL of 35% formaldehyde to preserve transcriptome information for scRNAseq. Cells were then fixed overnight at 4°C. For the wt barcode cells, the thawed cell pellet was resuspended in 4% formaldehyde in 1X PBS and incubated at room temperature for 15 minutes. After fixation, yeast-specific cell permeabilization and single-cell RNA sequencing preparations were performed using a yeast-specific application of split-pool based *in situ* transcriptomics sequencing (SPLiT-seq) as previously described in (Brettner et al. 2024), a caveat being that the wt barcode cells were alternately prepared with a commercially available Evercode WT Mini Parse kit starting at the reverse transcription step.

### Single-cell RNAseq data processing

#### Genome annotation

Sequencing reads were aligned to the R64 s288c genome from NCBI supplemented with added missing dubious ORFs from the iESR and transposable element sequences from the Saccharomyces Genome Database (SGD) (Cherry et al. 2012). We generated new fasta and GTF files with the 51 full length TE loci appended to the end. Internally annotated TE ORFs such as gag and pol genes (e.g., YLR035C-A), which lie within full-length TE loci, were masked to prevent redundant annotation and potential double counting of TE-derived reads. The FSA and GTF files are available on Open Science Framework (see below).

#### Transcriptome analysis

Transcriptome alignment, barcode assignment, and gene expression count analyses were performed as described in (Brettner et al. 2024). Reads were sorted by barcode using STARsolo (Kaminow et al. 2021). The yeast genome has significant sequence homology to itself due to a recent genome duplication (Wolfe 2015). Therefore, we utilized the uniform multimapping algorithm within STARsolo to keep reads that mapped to multiple locations in the genome. This means that multimapping reads were retained and assigned proportionally across all mapped locations, such that each location received a contribution equal to 1/n, where *n* is the number of mapped sites. This outputted gene-by-barcode matrices representing the detected gene counts for every cell. To remove barcodes that reflect empty cells and cells with low gene detection, we applied the “knee” detection filter previously described in (Brettner et al. 2024). These quality-thresholded gene-by-barcode matrices were then converted to Seurat Objects using the Seurat R packages Seurat (version 5.1.0) and SeuratObject (version 5.0.2). rRNA reads were bioinformatically removed before further analyses were performed (Hao et al. 2024). Computational work and visualizations were done with R (version 4.2.3).

Because SPLiT-seq relies on downstream recovery through reverse transcription, barcoding, library preparation, and sequencing, the number of successfully profiled cells varied across conditions and experiments. Differences in cell state during later growth phases may also contribute to the smaller number of profiled cells in these conditions apparent in **Fig 1D**.

Full analysis scripts are available on the Open Science Framework: https://osf.io/pcngh/files/osfstorage.

### Transposable element (TE) data processing

For analyses involving transposable elements (TEs), sequence homology posed an additional challenge. The *S. cerevisiae* genome contains multiple full-length copies of each Ty family, and loci within the same family can be highly similar in sequence. Consequently, many sequencing reads derived from a Ty1 element, for example, can align equally well to several different Ty1 loci. This ambiguity occurs primarily among loci within the same Ty family rather than across the five Ty families. As described above, STARsolo retained these multimapping reads and divided their contribution among the genomic locations to which they mapped.

This ambiguity is especially important in analyses where we define stress-response gene sets and then compare the number of genes in each set, overlap among sets, and the properties of the genes comprising each set (**Figs 1–3**). Treating every full-length TE locus as an independent “gene” would be misleading, because a read that can not distinguish among related TE copies can contribute fractional counts to many loci. A change in expression of a single Ty family could therefore appear as differences in dozens of separate “genes,” even though the data do not tell us that dozens of TE copies are independently changing. This could inflate the number of genes that differ between stress-specific gene sets, especially since the iESR does not include TEs. We therefore collapsed the 51 TE loci into five Ty-family features so that each family contributed at most one feature to these comparisons.

Reads were initially aligned using the 51 full-length TE loci described above, but counts assigned to loci belonging to the same family were subsequently summed. The individual TE loci were then replaced in the Seurat objects by five family-level features: Ty1, Ty2, Ty3, Ty4, and Ty5. Thus, each Ty family could contribute at most one feature to a stress-response gene set, rather than multiple highly homologous copies being counted as independent genes.

### Creating induced and repressed stress response gene lists (i.e., MSR, GSR, SSR, and FSR)

To determine the induced and repressed stress response genes, we compared gene expression of an unstressed population of cells to a stressed population of cells. For the MSR, we compared log-phase cells expressing YFPwt (**Fig 1A**; light purple) to those expressing YFPm4 (**Fig 1A**; dark purple). For the GSR we compared cells sampled early in glucose limitation (**Fig 1A**; lightest blue “T1”) to cells sampled late into glucose limitation (**Fig 1A**; darkest blue “T4”). For the SSR we compared YFPwt-expressing cells sampled early in sucrose limitation (**Fig 1A**; pink “T1”) to YFPwt-expressing cells sampled late in sucrose limitation (**Fig 1A**; red “T2”). For the FSR we compared cells sampled in exponential growth without fluconazole (**Fig 1A**; light green “No fluconazole”) to cells sampled at the same optical density (also in exponential growth) with 8 µg/mL fluconazole (**Fig 1A**; darkest green “High fluconazole”).

Differential expression analyses were performed on log normalized data. Because the number of sampled cells varied substantially among biological replicate flasks, an iterative balanced subsampling approach was implemented to minimize unequal contributions of individual flasks to the differential expression analysis while identifying genes whose expression changes were consistently observed across individual cells. For each comparison, cells from each biological replicate were randomly subsampled without replacement to match the number of cells in the smallest biological replicate, thereby ensuring balanced representation of each flask in every analysis. However, for the fluconazole treatment dataset, only a single biological replicate flask was available per fluconazole concentration. To conserve the detection of more uniformly differentially expressed genes, the cells from this flask were randomly subsampled to approximately 50% of the available cells for each iteration, producing subsamples while acknowledging the absence of biological replication.

Differential expression was performed on each balanced dataset using Seurat’s FindMarkers() function with the MAST statistical test. MAST employs a two-part hurdle model that separately models gene detection and expression magnitude, making it well suited for sparse single-cell RNA sequencing data with a high frequency of zero counts while maintaining sensitivity to genes exhibiting modest but consistent expression differences across many cells. It also allows for the consideration of latent, or covarying, variables in the time course glucose and sucrose data. For those two datasets, the latent variable was used to account for comparisons within the same flasks, just at different timepoints.

Balanced subsampling and differential expression analysis were repeated independently for 50 iterations comparing the stressed and unstressed conditions. For each iteration, log2 fold changes, raw *P* values, and Benjamini–Hochberg false discovery rate (FDR)-adjusted *P* values were calculated for every gene. Genes were considered robustly differentially expressed if they were significant following a slightly permissive Benjamini–Hochberg correction (FDR < 0.1) in at least 75% (38 of 50) of the subsampling iterations and exhibited a consistent direction of differential expression across significant iterations.

This resampling-based strategy reduces the influence of unequal cell numbers among biological replicates while prioritizing genes whose differential expression is reproducibly detected across independently balanced samplings of the underlying single-cell population.

While the MAST statistical test was used to create the induced and repressed gene lists for the stress conditions, the default Wilcoxon rank-sum test parameter was used in the calculations of the average log2 fold change (**Fig 2A**) and p-values of all genes for the volcano plots in **Fig S4** due to a large reduction in computational time.

For these analyses, counts from individual transposable element (TE) loci belonging to the same Ty family were summed to generate family-level features (e.g., Ty1 and Ty2). Thus, although reads were initially aligned to individual annotated TE loci, TE expression was analyzed at the family level when defining the MSR, GSR, SSR, and FSR gene lists.

### Calculating the percent transcriptome devoted to different gene sets (e.g., iESR)

The proportion of each cell’s transcriptome devoted to predefined gene sets was quantified using the PercentageFeatureSet() function in Seurat (v5). For each cell, the total unique counts assigned to genes within a specified gene set (e.g., iESR) were divided by the total unique counts detected for that cell and multiplied by 100, yielding the percentage of the cellular transcriptome represented by the gene set. These percentages were stored as cell-level metadata for downstream analyses and visualization. These data are plotted in **Fig 1B–1C** for the iESR and in **Supplemental Figs S2, S3, S5**, and **S6** for other gene sets.

### Ranking and classifying cells by their iESR (Fig 1C, D) or other gene sets

To assess whether cells could be classified as stressed or unstressed based on the degree of stress-response induction, we calculated for each cell the percentage of its transcriptome represented by predefined gene sets corresponding to induced or repressed stress responses. Cells were then ranked by these percentages for each stressed-versus-unstressed comparison (**Figs 1C, S2, S3, S5,** and **S6**). In the resulting ranked distributions, stressed and unstressed cells did not separate cleanly, indicating substantial overlap in stress-response magnitude between the two populations. To quantify classification performance, we ranked cells in ascending order by the percentage of transcriptome devoted to each induced stress-response gene set and assigned a threshold at the rank corresponding to the known number of unstressed cells. Cells below the threshold were classified as unstressed and cells above the threshold as stressed, and the percent of correctly classified cells was calculated for each group (**Figs 1D, S2C,** and **S3C**). For repressed stress-response gene sets, the classification was inverted, with cells below the threshold classified as stressed and cells above the threshold classified as unstressed (**Figs S5** and **S6**).

### Generating UMAPS for the stressed cell categories (Fig 2C)

Uniform Manifold Approximation and Projection (UMAP) visualizations were generated using Seurat (v5). For each stress condition, cells from the stressed population were merged across biological replicates. The percentage of each cell’s transcriptome represented by the relevant stress-response gene sets was calculated using PercentageFeatureSet() and stored as cell metadata prior to dimensionality reduction. For datasets containing multiple biological replicates, the RNA assay was split by replicate, normalized using SCTransform(), and integrated using Seurat’s IntegrateLayers() function with canonical correlation analysis (CCA) and SCT normalization. Principal component analysis (PCA) was performed on the SCT-normalized data, and UMAP embeddings were generated from the integrated CCA reduction using the first 10 principal components. For the fluconazole dataset, which contained a single biological replicate, no integration was performed; instead, PCA and UMAP were computed directly from the SCT-normalized data using the first 30 principal components. UMAPs were visualized using FeaturePlot(), with cells colored according to the percentage of their transcriptome represented by the indicated stress-response gene sets.

### Building Random Forest models and predicting cell provenance (Fig 3B and 3C)

Classifier-type Random Forest models (Breiman 2001) were built on log normalized transcriptomics data in R using the package randomForest (Breiman et al. 2002). Specifically, for every stress type dataset, cells were sorted into a stress (YFPm4, glucose T4, sucrose T2, or 8 ug/uL fluconazole) or an unstress (YFPwt, glucose T1, sucrose T1, or no fluconazole) category and given a label classification variable of “stress” or “unstress” to allow for model testing across datasets. Full models were then built for each dataset by comparing the classifier variable to the per cell gene expression for every annotated gene using largely the default parameters in the randomForest() function. Because our focus was on comparing model predictions across datasets rather than within a single dataset, we built each model with *all* cells from the given dataset instead of using training and testing data subsets. To calculate across model accuracy, we used the predict() function from the core R stats package to generate predictions from a given model using the other datasets as tests (e.g., misfolded protein, sucrose, and fluconazole datasets fed to the glucose model). We then used the confusionMatrix() function from the R package caret (Kuhn 2007), which calculates a cross-tabulation of observed and predicted categories to generate the accuracy statistics. These are the basis of the data displayed in **Fig 3B**. For **Fig 3C**, the same cross-dataset prediction procedure was repeated after withholding individual stress-response gene sets from the predictor variables. Each of the five induced gene sets (iESR, iMSR, iGSR, iSSR, and iFSR) and five repressed gene sets (rESR, rMSR, rGSR, rSSR, and rFSR) was removed separately from each of the four training datasets before retraining the Random Forest model, yielding 40 leave-one-gene-set-out models. Each retrained model was then used to predict stressed and unstressed cells from each of the other stress datasets, as described above. Prediction accuracy was compared with that of the corresponding full model to quantify the effect of withholding each gene set.

### Identifying genes differentially expressed in stressed cells with low iESR activation (Fig 4A)

To investigate the transcriptional profiles of cells in stressed conditions with low iESR expression, we pooled the data of cells from every stress condition (YFPm4, glucose T4, sucrose T2, and 8 ug/uL fluconazole) with the original TE annotation of all 51 full length loci into a single Seurat object and calculated the percent iESR using the PercentageFeatureSet() function as described above. We then sorted cells into the highest and lowest quartile of iESR percent expression and performed a differential expression analysis between the two groups using the FindMarkers() function with default parameters. The significantly differentially expressed genes in the low iESR group were then examined through a gene ontology analysis using Metascape (Zhou et al. 2019).

### Comparing the population expression of the iESR and transposable elements (Fig 4B)

To quantify the relationship between iESR and transposable element (TE) expression at the population level, pseudobulk transcriptomes were generated separately for each experimental condition. For each condition, raw UMI counts were summed across all cells to obtain total transcript counts for the iESR genes, Ty family transposable elements, and all expressed genes. The proportion of the transcriptome represented by the iESR and Ty transcripts was calculated by dividing the summed counts for each gene set by the total transcript counts for the condition, converting to counts per million (CPM), and applying a log10 transformation after adding a pseudocount of one [log10(CPM + 1)]. Separate linear regression models were then fit for each experimental dataset to evaluate the relationship between bulk Ty transcript abundance and bulk iESR expression across experimental conditions. Statistical analyses were performed using a Spearman rank correlation (**Fig 4D**) and the lm() function in R, and fitted regression lines were overlaid on scatter plots for visualization. The glucose diploid and haploid strains each have a single outlier as determined by the boxplot.stats() function. Those outliers remain on the **Fig 4B** plot and in the Spearman correlation calculation but were not used to fit the line to the two strains.

### Comparing the single-cell expression of the iESR and transposable elements (Figs 4C, D and S8, S9)

To quantify the relationship between stress-response activation and TE expression at the single-cell level, module scores were calculated for the induced Environmental Stress Response (iESR) (**Fig 4C, D**), induced Misfolding Stress Response (iMSR), induced Nutrient Starvation Responses to glucose (iGSR) and sucrose (iSSR), induced Fluconazole Stress Response (iFSR) (**Fig S9**), and the full length Ty transposable element loci using Seurat’s AddModuleScore() function. Module scores were calculated independently for each flask and sample following log-normalization of the raw UMI count matrix.

Because module scores may correlate with sequencing depth, each module score was tested for association with total cellular UMI counts using linear regression. When sequencing depth explained at least 10% of the variance in a module score (R² ≥ 0.10), residuals from the regression of module score against log-transformed total UMI counts were used in subsequent analyses; otherwise, the original module scores were retained. These values were used to create the single-cell scatter plots in **Fig 4C** and **S8**. Spearman rank correlations were then calculated between the residualized (or unadjusted) stress-response and TE module scores for each experimental condition.

To assess whether the observed correlations exceeded those expected by chance, two complementary null models were generated. In the first (“random read control”), 100 synthetic count matrices were generated by redistributing the observed cellular transcript counts according to the global gene abundance distribution while preserving the total read count of each cell. Module scores and sequencing-depth corrections were recalculated for each simulated dataset, and Spearman correlations between stress-response and TE module scores were used to estimate the correlation expected from transcript sampling alone. In the second (“random gene control”), 100 random gene sets matching the size of each stress-response gene set and the TE gene set were sampled without replacement from the expressed genes. Module scores were recalculated for each random gene set, corrected for sequencing depth as described above, and correlated with the corresponding observed module scores to estimate the background correlation expected from arbitrary gene selection. The median correlation across the 100 simulations was used as the representative value for each null model.

### Estimating probable full length Ty1 transcript expression and possible copy number control mechanisms (Fig S10)

The Ty1 family of TEs in *S. cerevisiae* possesses a well-characterized copy number control (CNC) mechanism that limits retrotransposition through interference with virus-like particle (VLP) assembly and maturation. The best-supported mechanism involves an internally initiated Ty1i transcript that originates within the GAG open reading frame and encodes p22, a trans-dominant truncated form of Gag. p22, and its proteolytically processed form p18, are incorporated into assembling VLPs, where they disrupt particle maturation and inhibit reverse transcription, thereby reducing Ty1 mobility (Saha et al. 2015; Tucker et al. 2015). Characterized Ty1 antisense (Ty1AS) RNAs that overlap the 5′ LTR and GAG have also been implicated in Ty1 regulation, although their mechanism remains less well resolved. They have been proposed to influence Ty1 transcription and/or VLP maturation, but their contribution to copy number control relative to the p22/p18 pathway remains uncertain (Berretta et al. 2008; Matsuda and Garfinkel 2009; Curcio et al. 2015). To establish if our Ty1 TE reads are actually dominated by CNC mechanisms and not full length Ty1 transcripts, we looked directly at the Ty1 alignments for properly barcoded sequencing reads that would make it into our downstream analyses.

SPLiT-seq combinatorial barcoding enabled reads derived from random-hexamer and poly-dT reverse-transcription primers to be analyzed separately. Primer identity was encoded in the round 1-specific barcode sequence, allowing reads from the same sequencing library to be assigned to either the random-hexamer or poly-dT primer class by using primer-specific barcode whitelist files during STARsolo alignment. STARsolo alignments were therefore run separately for the two primer-defined barcode sets, using the corresponding random-hexamer or poly-dT whitelist files.

Custom Ty1 annotations were generated separately for the random-hexamer and poly-dT alignments. For random-hexamer libraries, both the 5′ and 3′ LTRs were removed from Ty annotations. This was done because random priming should sample more broadly across the transcript, and including the highly similar 5′ and 3′ LTRs could artificially inflate the apparent 5′ Ty1 signal through multimapping between LTR copies. Removing both LTRs therefore restricted the random-hexamer analysis to the internal Ty1 sequence. For poly-dT libraries, the 5′ LTR was removed but the 3′ LTR was retained, because poly-dT priming is expected to enrich reads near the 3′ end of polyadenylated Ty1 transcripts, making the 3′ LTR a biologically relevant region for this primer class.

Ty1 annotations were further divided into “true” and “ambiguous” regions relative to the internally initiated Ty1 CNC/p22 transcript. The upstream gag region before the p22/Ty1i start was treated as true full-length Ty1 signal, because reads overlapping this region are incompatible with being derived solely from the CNC transcript. The downstream region beginning at the p22/Ty1i start was treated as ambiguous, because reads in this region could derive either from full-length Ty1 RNA or from the CNC/p22 transcript. A separate antisense bin was defined over the known Ty1 antisense RNA region.

To estimate the fraction of Ty1 signal attributable to true, ambiguous, and antisense regions, coordinate-sorted BAM files were parsed directly. Only Ty1 alignments from reads with valid STARsolo barcode evidence were retained. To avoid counting the same sequencing read multiple times, alignments with the same read name were grouped together before classification, so each read contributed a maximum of one count despite possible multimapping among Ty1 copies. In the final conservative classification, any forward-strand Ty1 read with an aligned span overlapping the upstream true gag region was assigned to the true Ty1 bin. Reads were assigned to the ambiguous bin only if their forward-strand alignment was fully contained within the post-p22 region. Reverse-strand reads overlapping the known Ty1 antisense interval were assigned to the antisense bin, and reads outside these criteria were classified as other forward or other reverse. Each read was counted once in the highest-priority applicable category.

## Supplemental Table Legends

**Supplemental Table 1:** A table with all gene sets used in this study. iESR and rESR genes are from (Gasch et al. 2000; Brauer et al. 2008; Ho and Gasch 2015; Costa-Mattioli and Walter 2020; Chen et al. 2023).

**Supplemental Table 2:** A table with Gene Ontology (GO) results for each induced stress gene set on a different tab (iMSR, iGSR, iSSR, and iFSR) (Zhou et al. 2019).

## Data Availability Statement

Data processing and analysis scripts, as well as processed alignment data can be found on the Open Science Framework: https://osf.io/pcngh/files/osfstorage. Both raw fastq files and alignment data for the glucose stress experiment and the first misfolding/sucrose stress experiment can be accessed through NCBI GEO GSE251966 (data used in this study pertains to samples GSM7990658 and GSM7990659). Both raw fastq files and alignment data for the second misfolding/sucrose stress experiment and the fluconazole stress experiment can be accessed through NCBI GEO GSE309715. We also provide raw sequencing reads as fastq files, along with raw, unfiltered barcode (cell)-by-gene count matrices that were created after first aligning all sequenced reads to the appropriate genomes using STARsolo and assigning each read according to its corresponding barcode. All subsequent analyses performed in this publication can be reproduced directly with these data.

## Acknowledgments

Funding for this work was provided by a National Institutes of Health grant R35GM133674 (to KGS), an Alfred P Sloan Research Fellowship in Computational and Molecular Evolutionary Biology grant FG-2021-15705 (to KGS), and a National Science Foundation Biological Integration Institution grant 2119963 (to KGS). We would also like to thank the members of the ASU Biodesign Center for Mechanisms of Evolution and the Geiler-Samerotte Lab for their support and feedback.

## Competing Interests

The authors have declared that no competing interests exist.

## Supplemental Figures

**Fig S1:**
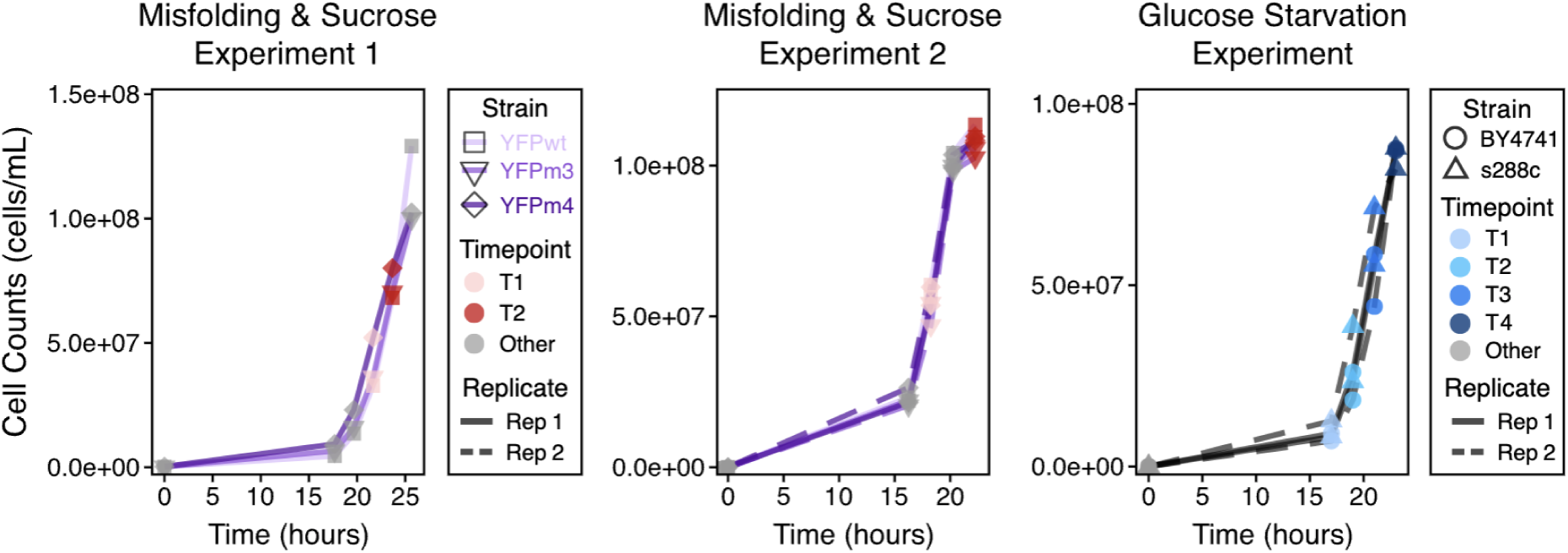
Growth data showing cell counts and sampling. Gray points indicate timepoints when cells were counted but not sampled for scRNAseq. Colored points represent timepoints at which scRNAseq was performed. In the leftmost two graphs, misfolding stress was surveyed during timepoint 1 (pink) by comparing transcriptomes across strains with different levels of misfolding. Sucrose stress was assessed by comparing, within each strain, transcriptomes from the YFPwt strain at timepoint 1 (pink) versus timepoint 2 (red). In the rightmost graph, glucose stress was surveyed by comparing transcriptomes from timepoint 1 (lightest blue) to those from timepoint 4 (darkest blue). Both of the rightmost graphs include two flasks (replicates) pers strain. The leftmost graph includes a single flask per strain.

**Fig S2:**
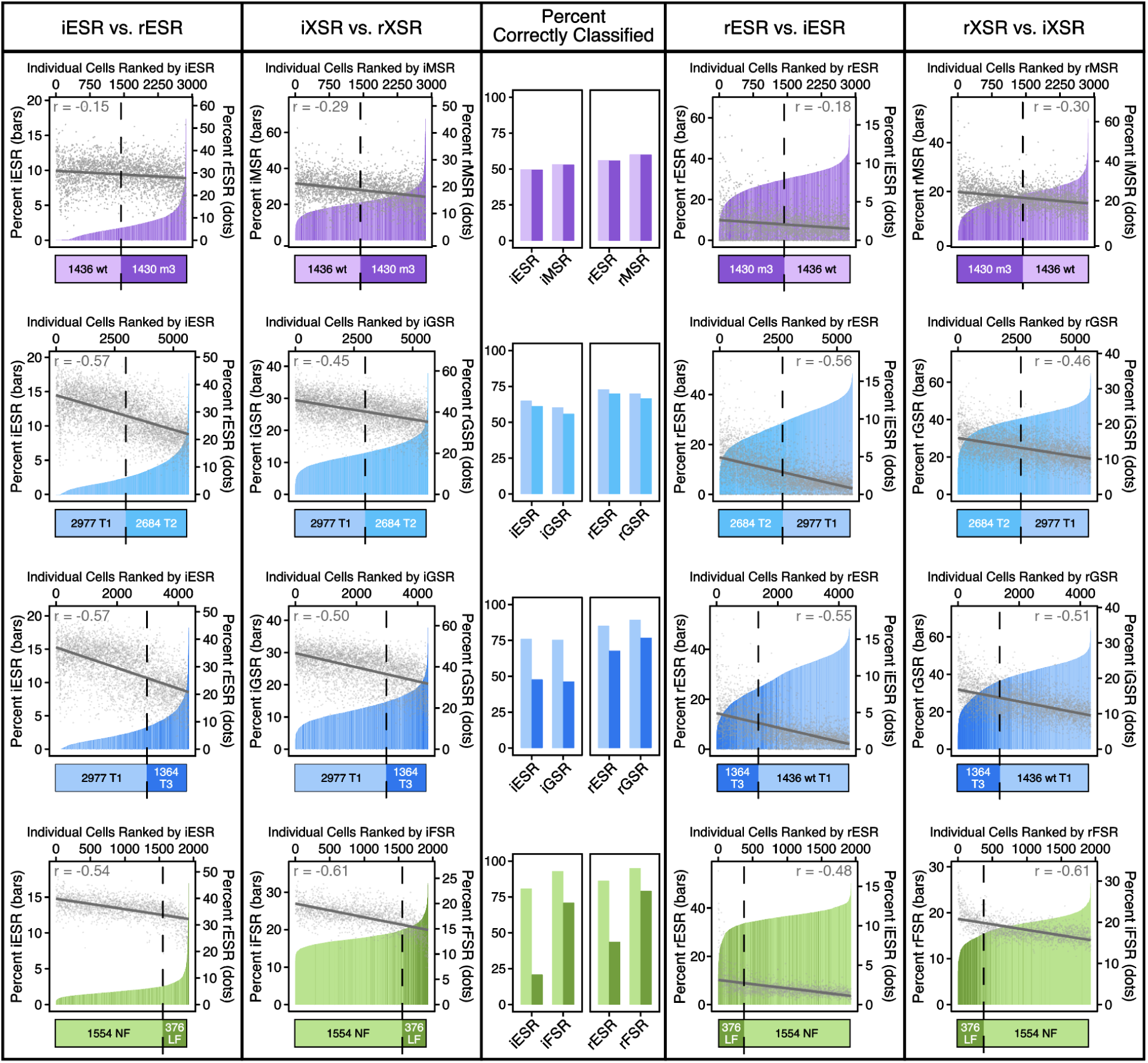
Single-cell heterogeneity across induced stress responses in additional milder stress comparisons. **Rows:** (1) YFPwt and YFPm3, (2) T1 and T2 (glucose), (3) T1 and T3 (glucose), (4) No fluconazole and Low fluconazole. **Leftmost column:** Cells are ordered by the percent of their transcriptome dedicated to the iESR (bars). These percents are calculated using the Seurat function PercentFeatureSet(). An inverse correlation (Pearson coefficient shown on graph) is observed between the (induced) iESR and (repressed) rESR (points). Linear trend lines are also negative. The vertical dashed line signifies the true number of cells from the unstressed condition; horizontal bars indicate the true number of unstressed and stressed cells in each plot. **Second column:** Similar to leftmost column except cells are ordered by the percent of their transcriptome dedicated to corresponding induced stress responses (bars). **Middle column:** Bars display the number of correctly classified unstressed or stressed cells based on the rank order of their iESR, rESR, or corresponding induced or repressed stress response. **Fourth column:** Similar to the leftmost column except cells are ordered by the percent of their transcriptome dedicated to the rESR (bars) and the points depict each cell’s level of the iESR. **Rightmost column:** Similar to the fourth column except cells are ordered by the percent of their transcriptome dedicated to the corresponding repressed stress response (bars).

**Fig S3:**
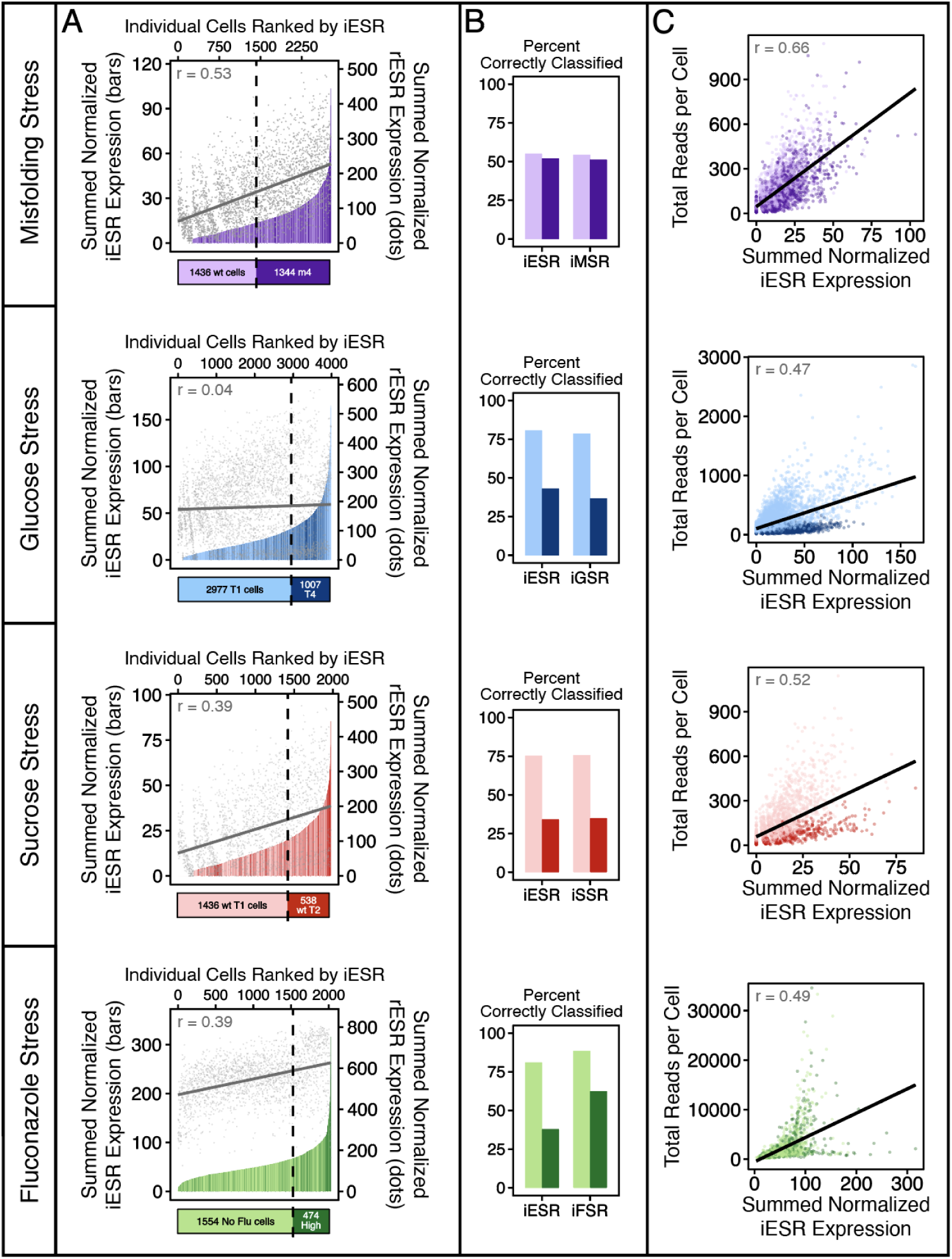
Summed normalized count of iESR genes scales with total reads per cell and is therefore not useful for predicting cell provenance. **(A)** These plots are similar to those in **Fig 1C** but instead of using the Seurat function PercentFeatureSet(), we quantify each cell’s iESR level by using summed normalized counts. These summed counts do not cleanly distinguish cells that came from a stressful or unstressful condition as evidenced by the intermixing of darker colored bars and lighter colored bars along the horizontal axis of each plot. **(B)** Bars display the number of correctly classified unstressed or stressed cells based on the rank order of their iESR, rESR, or corresponding induced or repressed stress response. **(C)** Every point represents a single cell. Total reads per each cell (vertical axis) positively correlates (Pearson coefficient shown on graph) with the summed normalized iESR reads per cell (horizontal axis).

**Fig S4:**
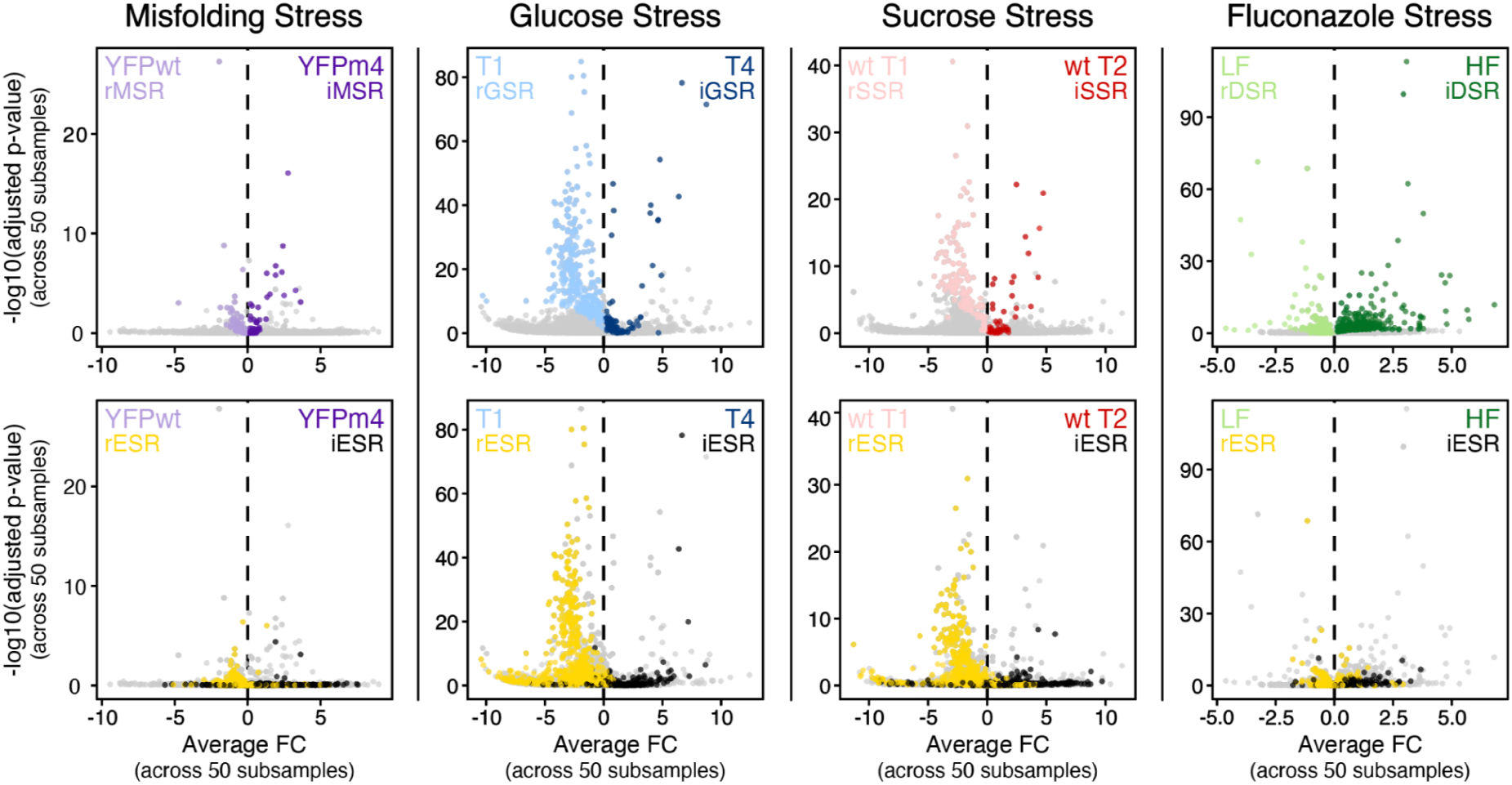
Volcano plots showing fold change of genes comprising each stress response. For each of the four stress types (misfolding, glucose, sucrose, and fluconazole), we subsampled cells from the stressed and unstressed conditions 50 times each, calculated the average fold change in expression for each gene across subsamples, and then assessed whether that fold change was significant based on the variation across subsamples (see Methods). When multiple levels of stress were surveyed, for example YFPm3 and YFPm4, we focused on the strongest stress level for this comparison, as denoted on each plot. In the top row, the genes that passed our significance threshold are colored. In the bottom row, the genes comprising the rESR (yellow) and iESR (black) are colored (all other genes are gray). A key observation is that the genes comprising the iESR (black) remain low on the vertical axis, and are sometimes found on the left rather than the right side of the vertical line, and are thus not significantly upregulated in stress using our metric.

**Fig S5:**
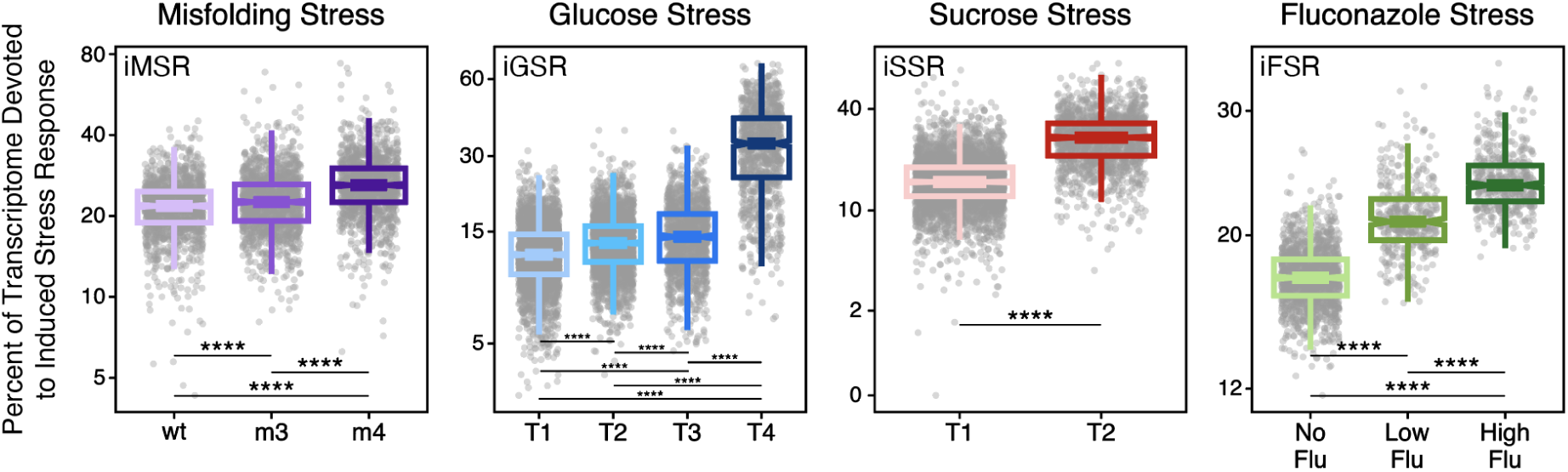
Percent of transcriptome devoted to induced stress responses increases as stressor increases. The percent of the transcriptome devoted to stress responses defined using our single-cell-aware approach (iMSR, iGSR, iSSR, and iFSR). Every dot represents a single cell; median and quartiles are shown across cells pertaining to a given strain (misfolding stress), timepoint (nutrient stress), or fluconazole level. **** indicates *p* < 0.0001 and *** indicates *p* < 0.001 with Wilcoxon rank-sum test.

**Fig S6:**
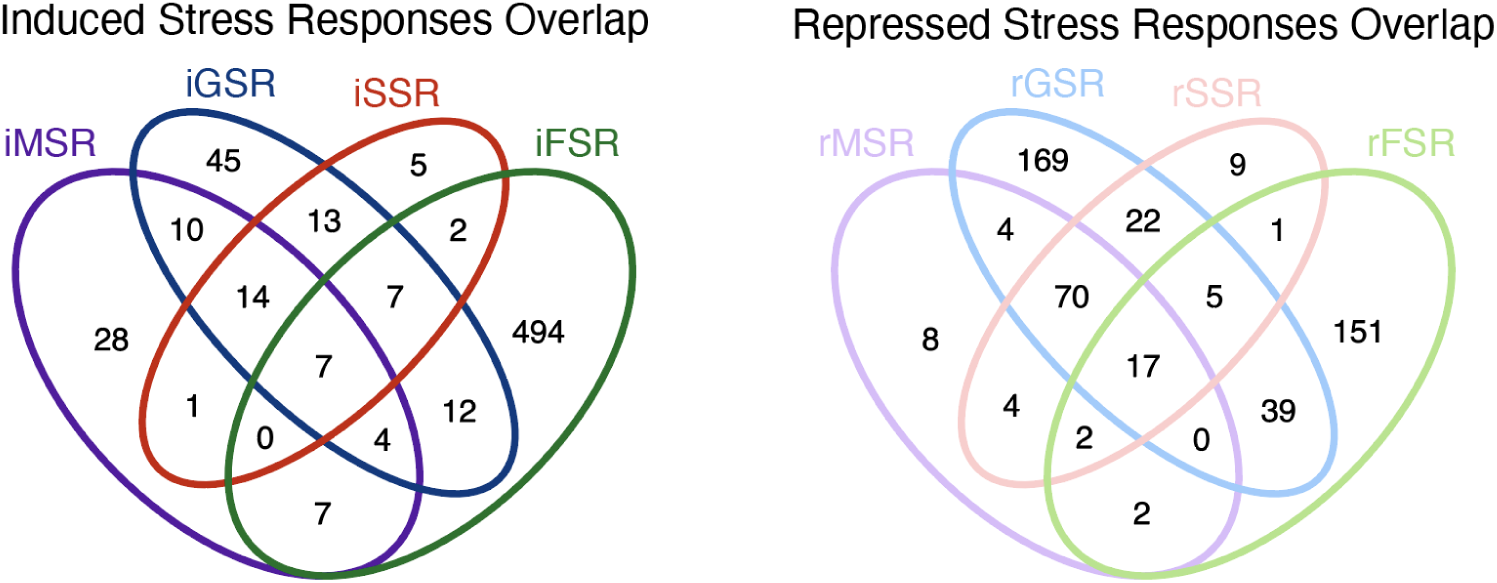
Venn diagrams showing the overlap between the genes that comprise the induced and repressed stress responses defined using our single-cell-aware approach. The numbers inside the Venn diagrams correspond to the number of genes.

**Fig S7:**
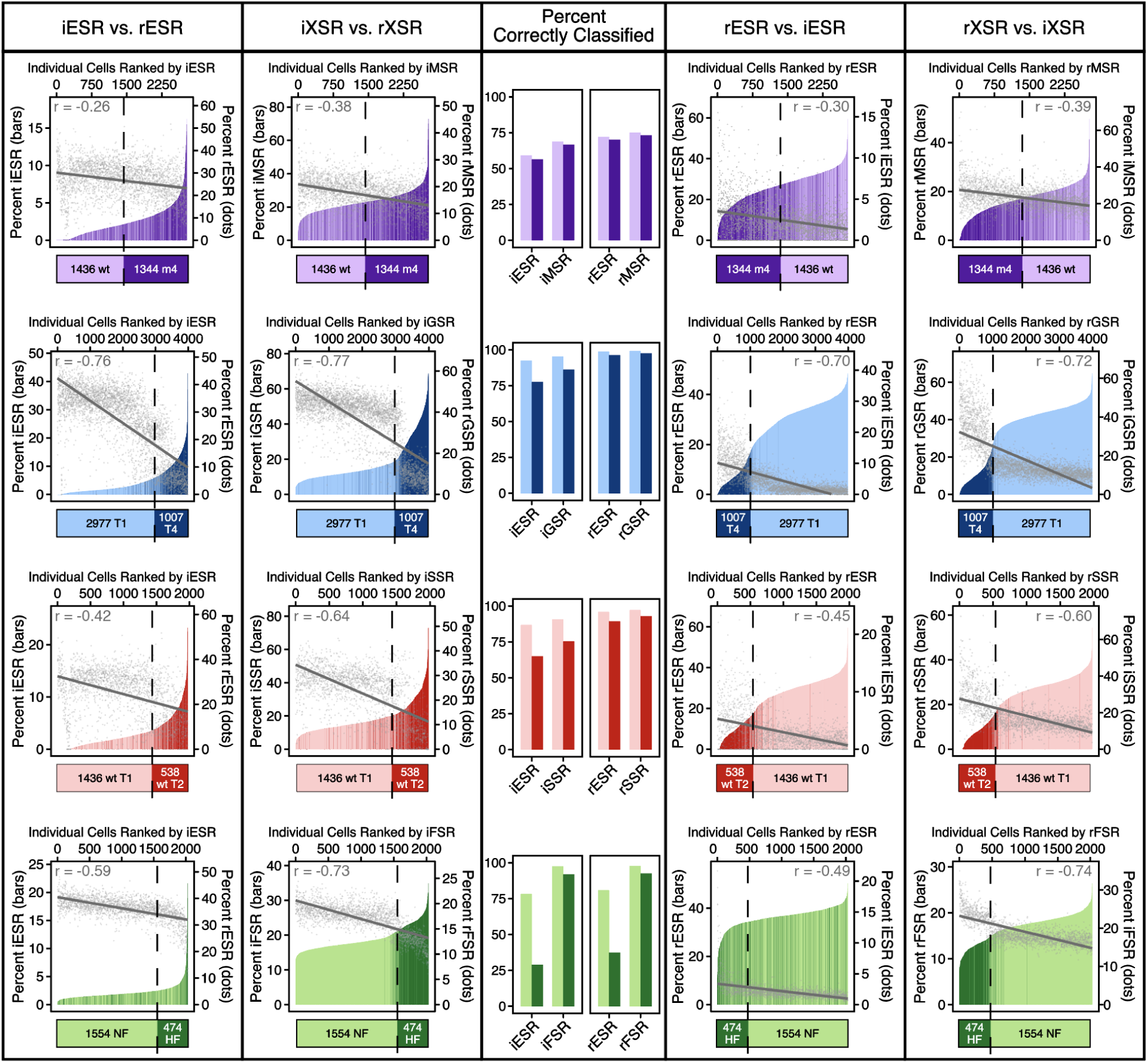
Single-cell heterogeneity across induced stress responses; comparison of threshold classifiers based on different induced and repressed stress responses. **Rows:** (1) YFPwt and YFPm4, (2) T1 and T4 (glucose), (3) T1 and T2 (sucrose), (4) No fluconazole and High fluconazole. **Leftmost column:** This column is reproduced from **Fig 1C** to allow for comparison with the other columns. Cells are ordered by the percent of their transcriptome dedicated to the iESR (bars). These percents are calculated using the Seurat function PercentFeatureSet(). An inverse correlation (Pearson coefficient and linear regression shown on graph) is observed between the (induced) iESR and (repressed) rESR (points). Linear trend lines are also negative. The vertical dashed line signifies the true number of cells from the unstressed condition; horizontal bars indicate the true number of unstressed and stressed cells in each plot. **Second column:** Similar to leftmost column except cells are ordered by the percent of their transcriptome dedicated to corresponding induced stress responses defined using our single-cell aware approach (bars). **Middle column:** Bars display the number of correctly classified unstressed or stressed cells based on the rank order of their iESR, rESR, or corresponding induced or repressed stress response. **Fourth column:** Similar to the leftmost column except cells are ordered by the percent of their transcriptome dedicated to the rESR (bars) and the points depict each cell’s level of the iESR. **Rightmost column:** Similar to the fourth column except cells are ordered by the percent of their transcriptome dedicated to the corresponding repressed stress response defined using our single-cell-aware approach (bars).

**Fig S8:**
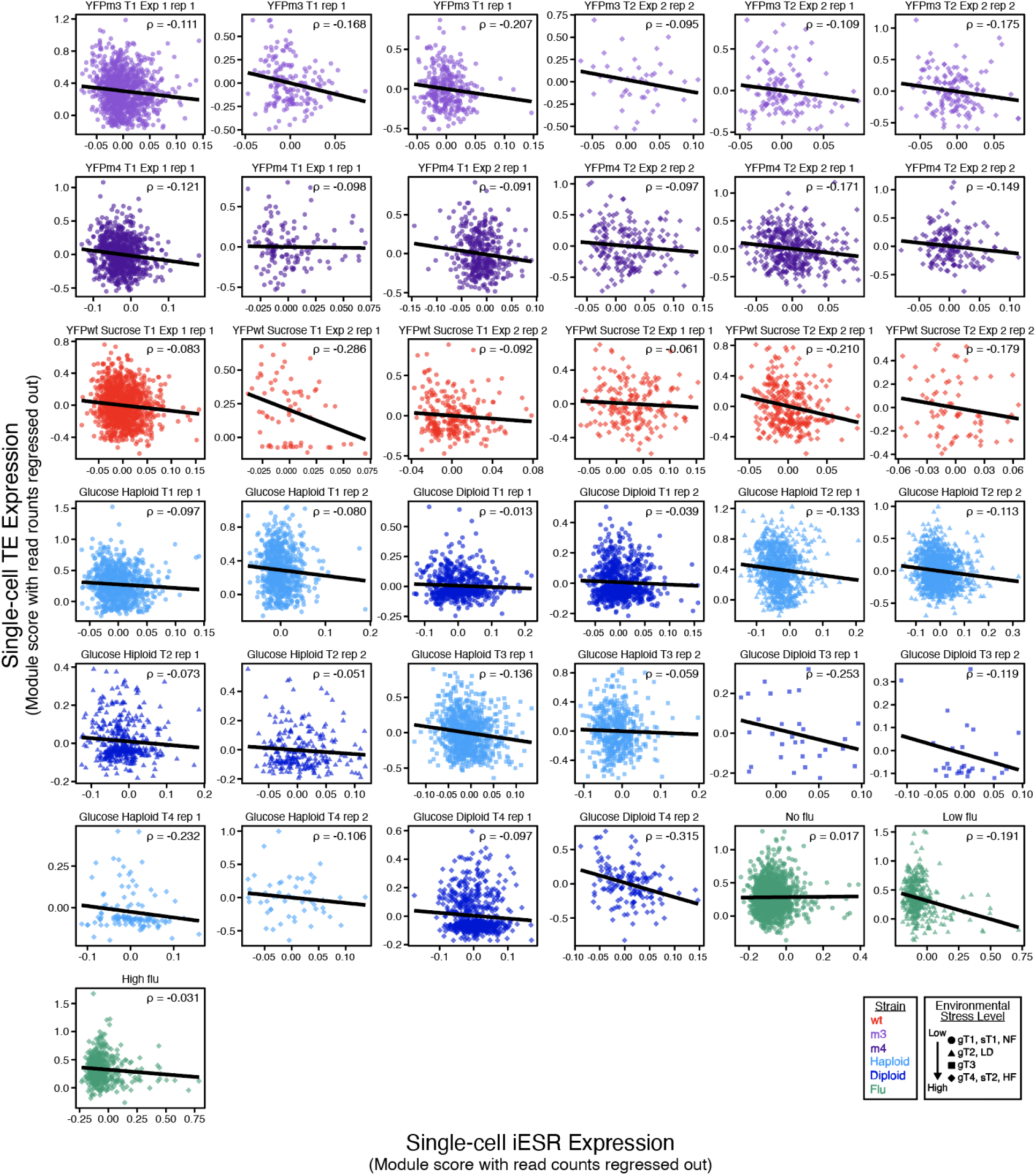
TEs and the iESR anticorrelate on the single-cell level in all 37 samples. The iESR module score (horizontal axis) and TEs module score (vertical axis) are plotted for each individual cell (a point on the graph) for all 37 samples, colored as in **Fig 4B**. Every sample represents an independent flask of cells. Trend lines indicate an anticorrelation or near-zero correlation for all plots. Spearman correlations are included in **Fig 4D** “iESR vs. TEs.”

**Fig S9:**
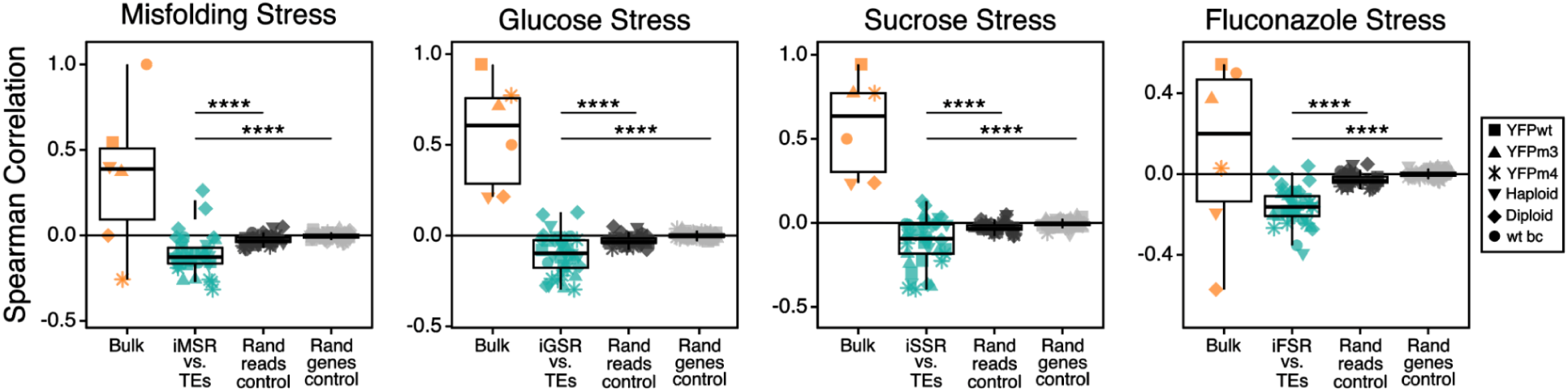
TEs and all stress-specific induced responses anticorrelate on the single-cell level. Similar to **Fig 4D** but for the stress responses defined here using our single-cell-aware approach.

**Fig S10:**
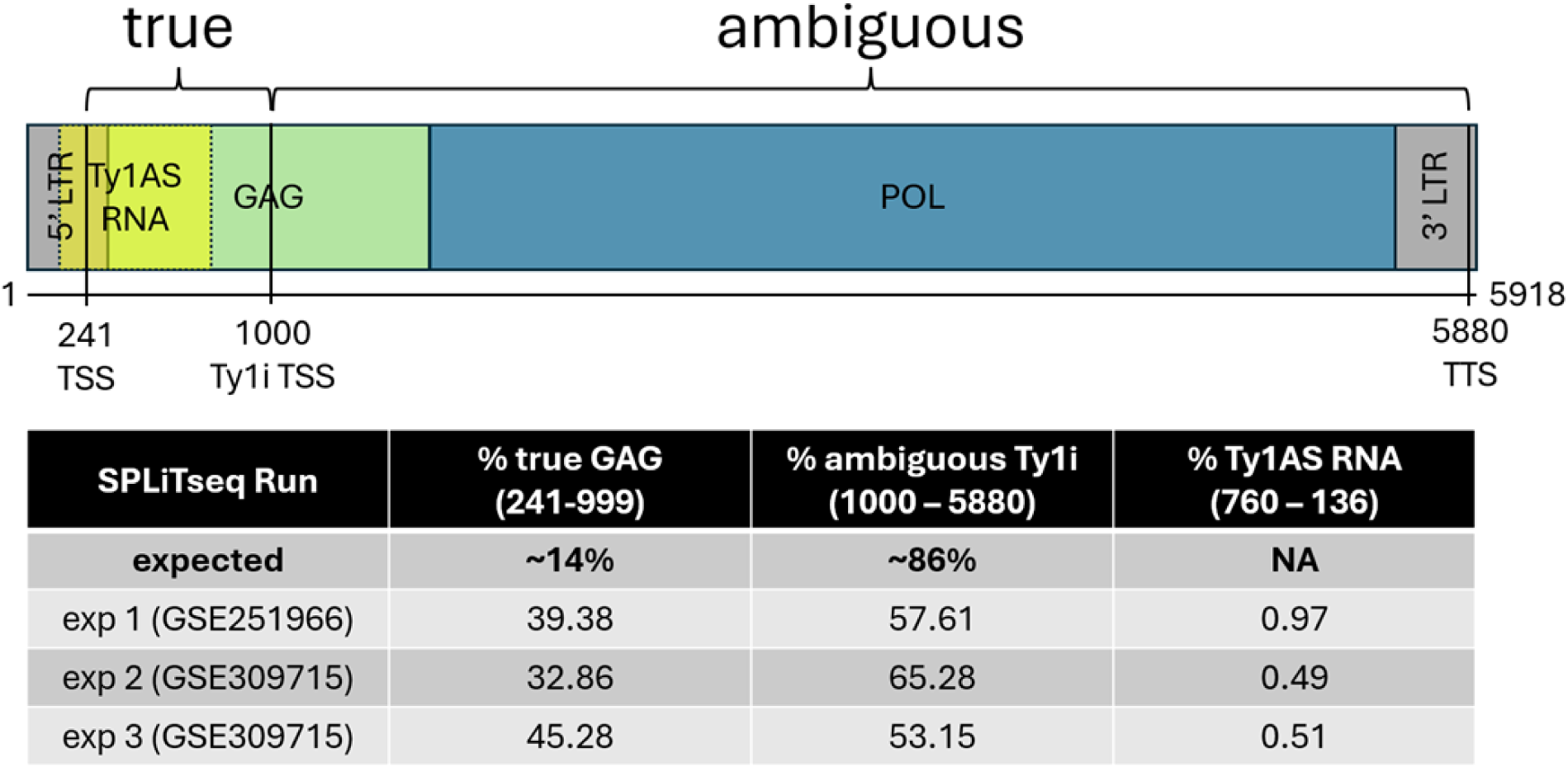
TE Ty1 family copy number control (CNC) transcripts are unlikely to dominate Ty1 reads. The canonical Ty1 transcript is initiated at approximately position 241 from the 5′ LTR promoter and encodes the Gag and Pol proteins required for retrotransposition. CNC is mediated by an internally initiated Ty1i transcript originating within the GAG open reading frame at approximately position 1000. Translation of the Ty1i transcript produces the trans-dominant Gag-derived restriction factor p22, which is processed to p18 during virus-like particle (VLP) maturation. p22/p18 disrupts VLP assembly and maturation, thereby inhibiting reverse transcription and reducing Ty1 retrotransposition (Saha et al. 2015; Tucker et al. 2015). Ty1 antisense (Ty1AS) RNAs overlapping the 5′ LTR and GAG have also been described and have been implicated in Ty1 regulation, although their mechanism and contribution to copy number control remain unresolved (Matsuda and Garfinkel 2009; Curcio et al. 2015). Ty1 aligned reads were assigned to regional bins and counted: true Ty1 if they overlapped pre-Ty1i Gag sequence, ambiguous if fully contained downstream of the Ty1i start, and antisense if mapping to the known reverse-strand Ty1 antisense region (yellow) (see Methods). Given a conservative uniform distribution of reverse-transcription priming sites across full-length Ty1 RNA, approximately 14% of barcoded reads would be expected to map to the pre-Ty1i true Gag region based on its relative length. However, all sequencing runs showed more than twice this expected fraction. This enrichment could reflect several non-mutually exclusive factors, including increased priming near an A-rich sequence close to the Ty1i/p22 start region at position 1065. Regardless of mechanism, the high fraction of reads mapping to the 5′ full-length Ty1-specific region suggests that an overabundant representation of the p22/CNC transcript in our data is unlikely. Similarly, reads mapping to the known Ty1 antisense region comprised less than 1% of Ty1 reads. Together, these results indicate that the Ty1 signal captured in our sequenced libraries predominantly reflects full-length Ty1-associated transcripts rather than either copy number control mechanism.

## Notes

### Competing Interest Statement

The authors have declared no competing interest.

### Summary of Updates

Adding an independent scRNA-seq dataset representing a distinct class of stress Repeating key analyses using additional analytical approaches and showing that they support the same biological conclusions Substantially revising the analyses underlying all four main figures Adding numerous new controls and validation analyses to test alternative technical explanations Expanding methodological detail, reproducibility information, and engagement with prior single-cell literature Reorganizing and rewriting the manuscript throughout to reflect the revised conceptual framework

